# Getting Personal: Brain Decoding of Spontaneous Thought Using Personal Narratives

**DOI:** 10.1101/2023.05.12.540141

**Authors:** Hong Ji Kim, Byeol Kim Lux, Emily S. Finn, Choong-Wan Woo

## Abstract

The contents of spontaneous thought and their dynamics are important factors for one’s personality traits and mental health. However, they are difficult to assess because spontaneous thought occurs voluntarily without conscious constraints. Here, we aimed to decode two important content dimensions of spontaneous thought—self-relevance and valence—directly from functional Magnetic Resonance Imaging (fMRI) signals. To train brain decoders, we induced a wide range of levels of self-relevance and emotional valence using individually generated personal stories as well as stories written by others to mimic narrative-like spontaneous thoughts (*n* = 49). We then tested the brain decoders on two resting-state fMRI datasets (*n* = 49 and 90) with and without intermittent thought sampling, achieving significant predictions. The default mode and ventral attention networks were important contributors to the predictions. Overall, this study paves the way for the brain decoding of spontaneous thought and its use for clinical applications.

## Introduction

Our mind never rests. Even during rest or sleep, our mind spontaneously wanders from the past to the future and from one concept to another^1–3^. Spontaneous thoughts may seem random, but they often involve topics that are emotionally charged, central to self-identity, and related to internal desires and goals^4, 5^. The contents and their dynamics of spontaneous thought are known to be important predictors of cognitive and affective traits (e.g., ruminative or internalizing cognitive styles)^2, 6–8^ and disrupted brain processes, providing potential as cognitive and behavioral markers for mental and neurologic disorders, such as depression, anxiety, and Alzheimer’s disease^9–11^. However, the assessment of one’s spontaneous thought is challenging given that it occurs freely with minimal conscious constraints^12^. In addition, the act of paying attention to spontaneous thought can change the nature of spontaneous thought itself, also known as the Heisenberg effect^13^. For these reasons, measuring some aspects of spontaneous thought directly from brain activity, e.g., functional Magnetic Resonance Imaging (fMRI) signals, would be useful. The aim of this study is to examine whether we could decode two key content dimensions of spontaneous thought—self-relevance and valence^6^—by developing functional neuroimaging-based predictive models. This can be viewed as an effort for brain-based daydream decoding, similar to dream decoding^14^, but specifically targeting affective aspects of spontaneous thought.

To effectively induce brain representations that resemble those of spontaneous thought, we chose to use narratives as stimuli. Recent studies have suggested that spontaneous thought is experienced in the form of images or words^15^, particularly as deeply processed imagery and concepts, e.g., narratives^16^ and scene construction ^17^. Though narratives cannot capture all aspects of spontaneous thought, narratives share key elements with spontaneous thought, such as rich semantic information and their temporally unfolding nature^18^. In addition, narratives have been successfully used in fMRI experiments to study semantic processing in the brain^19–23^. Thus, narratives provide promising candidate materials for the study of brain representations of spontaneous thought.

However, the contents of spontaneous thought have an important characteristic that narratives created by others (e.g., experimenters) are lacking, which is personal relevance (or self-relevance)^7, 24, 25^. This motivated us to use personal narratives in our experiment. It is well-known that spontaneous thought is most commonly about one’s personal life, such as current personal concerns, past memories, and future plans^5, 7, 26–28^, suggesting that self-relevant thought contents are major building blocks of spontaneous thought. For example, a recent paper proposed that episodic memory serves as a foundation of spontaneous thought and provides a scaffolding for semantic memory to generate thought contents^29^. In addition, self-referential information recruits brain systems distinct from those for non-self-referential information^30–33^, underscoring the importance of using self-relevant stimuli to study brain representations of spontaneous thought. We, thus, hypothesized that personal narratives would be able to induce brain representations close to those of spontaneous thought. Here, we performed one-on-one interviews with participants to create individually unique stimuli based on personal narratives, which were used in the fMRI experiment to elicit self-relevant thoughts and emotions.

Among multiple dimensions of spontaneous thought^6, 7, 15^, here we focus on two content dimensions—self-relevance and valence. We chose to focus on these two dimensions mainly because dimensionality reduction analyses conducted in previous studies showed that self-relevance and valence were among the most central dimensions that can serve as summaries of other content dimensions^6, 7^. In addition, these two variables are likely to be among the fundamental dimensions of human cognition and emotions given that self-relevance and valence convey information crucial for survival, such as what to avoid (i.e., negative valence, high self-relevance), what to approach (i.e., positive valence, high self-relevance), or what to ignore (i.e., low self-relevance). In terms of the brain systems, self-relevance and valence can be linked to the brain networks related to valuation (i.e., is it good or bad?) and context-dependent salience detection (i.e., is it relevant to me?), such as the default mode, limbic, and ventral attention networks^34–36^. Note that, however, these two dimensions are only a small fraction of components that constitute spontaneous thoughts, and therefore consider this study as the first step to the decoding of the rich contents of spontaneous thought.

As shown in **Fig. 1a**, the major research goals of the current study include 1) developing fMRI multivariate pattern-based predictive models of self-relevance and valence using data from the story-reading task, 2) comparing and interpreting the newly developed predictive models of self-relevance and valence, and 3) testing the predictive models on the resting-state fMRI data with and without intermittent thought-sampling probes. To this end, we conducted an fMRI experiment (*n* = 49) while participants underwent the story-reading and thought-sampling tasks.

**Figure 1.**
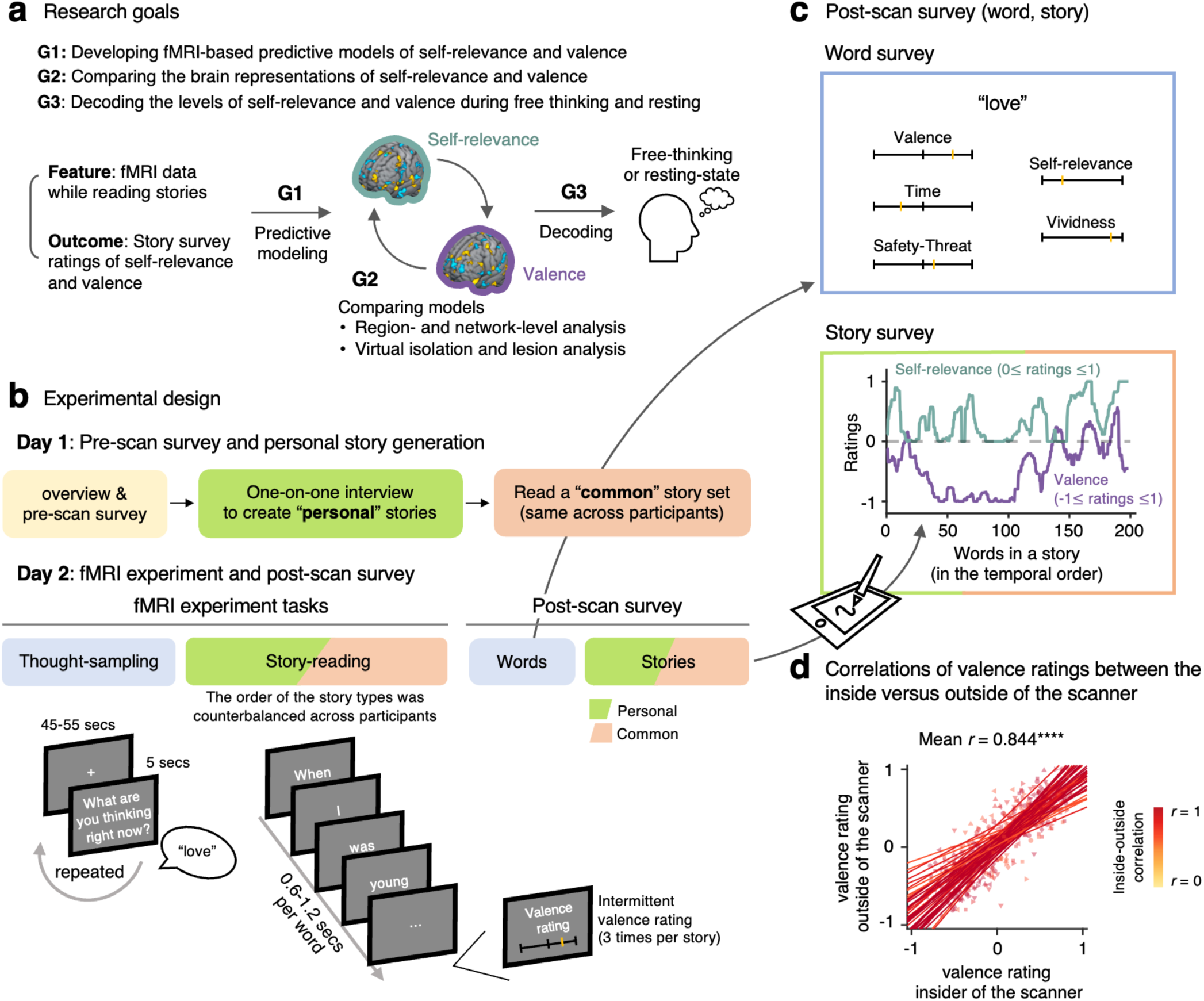
Research goals and experimental design. **a,** This study has three research goals, each corresponding to different analysis steps. The first goal is to build predictive models, the second goal is to interpret the model features that contribute most to prediction, and the third goal is to test the models on independent data. **b,** The experiment was conducted over two days separated by an interval of approximately one-week (mean = 7.3 days). On Day 1, we conducted a pre-scan survey and one-on-one interviews to create personal story stimuli. On Day 2, we conducted an fMRI experiment that consisted of 5 story-reading runs and 2 thought-sampling runs. During the story-reading runs, we asked participants to intermittently rate the current emotional valence (three times per story). For details of the experimental procedure, please see **Methods**. **c**, After the scan, participants underwent the post-scan survey for thought-sampling responses (i.e., words or phrases) and stories. For each thought-sampling response, we asked participants to rate the response on five content dimensions, including valence, self-relevance, and other dimensions. For the story survey, we asked participants to listen to the stories again while continuously rating them on the content dimensions of valence and self-relevance using a tablet pen. The plot shows example story ratings, in which the green line indicates self-relevance ratings, while the purple line indicates valence ratings. **d,** The valence ratings between the inside and the outside of the scanner showed high within-individual correlations, mean *r* = 0.844, *p* < 2.220e-16, two-tailed, bootstrap tests with 10,000 iterations.

In the story-reading task, participants were asked to read their own stories or stories made by others to induce a wide range of levels of self-relevance and valence. In the thought-sampling task, participants were asked to think freely and intermittently report a few words that represented their current thought. After fMRI scans, participants provided self-relevance and valence ratings for the stories and words from the story-reading and thought-sampling tasks. With the fMRI data from the story-reading task, we developed fMRI multivariate pattern-based decoding models of self-relevance and valence that showed significant predictions in the leave-one-subject-out (LOSO) cross-validation. We then identified important contributors to the prediction of both models using the virtual lesion and isolation analysis methods. Finally, we applied these models to decode self-relevance and valence scores during the thought sampling task (*n* = 49) and resting state (*n* = 90).

## Results

### Experimental overview and post-scan survey

**Fig. 1b** shows the experimental design of the current study, which we briefly describe here (for the details of the experimental procedure, please see **Methods**). On Day 1, we conducted a one-on-one interview with participants to create personal stories to use as stimuli in the fMRI experiment. On Day 2, participants underwent the story-reading and thought-sampling tasks in the scanner. During the story-reading task, participants read four “personal” stories, which were created for each participant, and six “common” stories, which were the same across participants. While reading the stories, participants were intermittently asked to provide their valence ratings (i.e., three times per story). In the thought-sampling task, we asked participants to think freely and verbally report what was in their mind with a few words intermittently (every 50.7 ± 5.6 [mean ± SD] seconds). After the scan, we conducted a post-scan survey on words and stories (**Fig. 1c**). For the word survey, participants rated the words they generated during the thought-sampling task using a multi-dimensional content scale (see **Methods** for details), and for the story survey, participants read the stories again and rated their perceived levels of self-relevance and valence using continuous ratings (see **Supplementary Fig. 1** for example and group average ratings). To ensure that the post-scan survey results reflected the in-scanner experience, we compared the intermittent valence ratings from the in-scanner story-reading task with the valence ratings from the post-scan survey. As shown in **Fig. 1d**, the in-scanner vs. post-scan valence ratings were highly correlated (mean *r* = 0.844, z = 51.71, *p* < 2.220e-16, two-tailed, bootstrap tests with 10,000 iterations), suggesting that the post-scan survey ratings reflected the in-scanner experience well.

### Developing predictive models of self-relevance and valence

To achieve the first research goal, i.e., developing fMRI-based predictive models of self-relevance and valence (**Fig. 1a**), we trained fMRI multivariate pattern-based predictive models using the fMRI data from the story-reading task. To effectively disentangle fMRI patterns for self-relevance and valence, we concatenated and quantized the data based on quintiles of the two dimensions, constructing 25 averaged images per participant (i.e., 5 levels of self-relevance × 5 levels of valence; for details of the data quantization and distribution of ratings, see **Supplementary Figs. 2-3**). We then trained predictive models of self-relevance and valence using principal component regression (PCR)^37^ with leave-one-subject-out cross-validation (LOSO-CV) and random-split cross-validation (RS-CV)^38, 39^. The models showed significant cross-validated prediction performances—prediction-outcome correlations for self-relevance, with LOSO-CV, mean *r* = 0.322, *z* = 9.204, *p* < 2.220e-16, mean squared error (mse) = 0.148, two-tailed, bootstrap tests with 10,000 iterations, with RS-CV, mean *r* = 0.332, mse = 0.144 (**Fig. 2a**); for valence, with LOSO-CV, mean *r* = 0.205, *z* = 6.235, *p* = 4.511e-10, mse = 0.454, two-tailed, bootstrap tests with 10,000 iterations, with RS-CV, mean *r* = 0.179, mse = 0.458 (**Fig. 2b**). Permutation tests with 1,000 iterations and LOSO-CV also provided significant performances against null models (both *p* = 0.0010; **Supplementary Fig. 4**).

**Figure 2.**
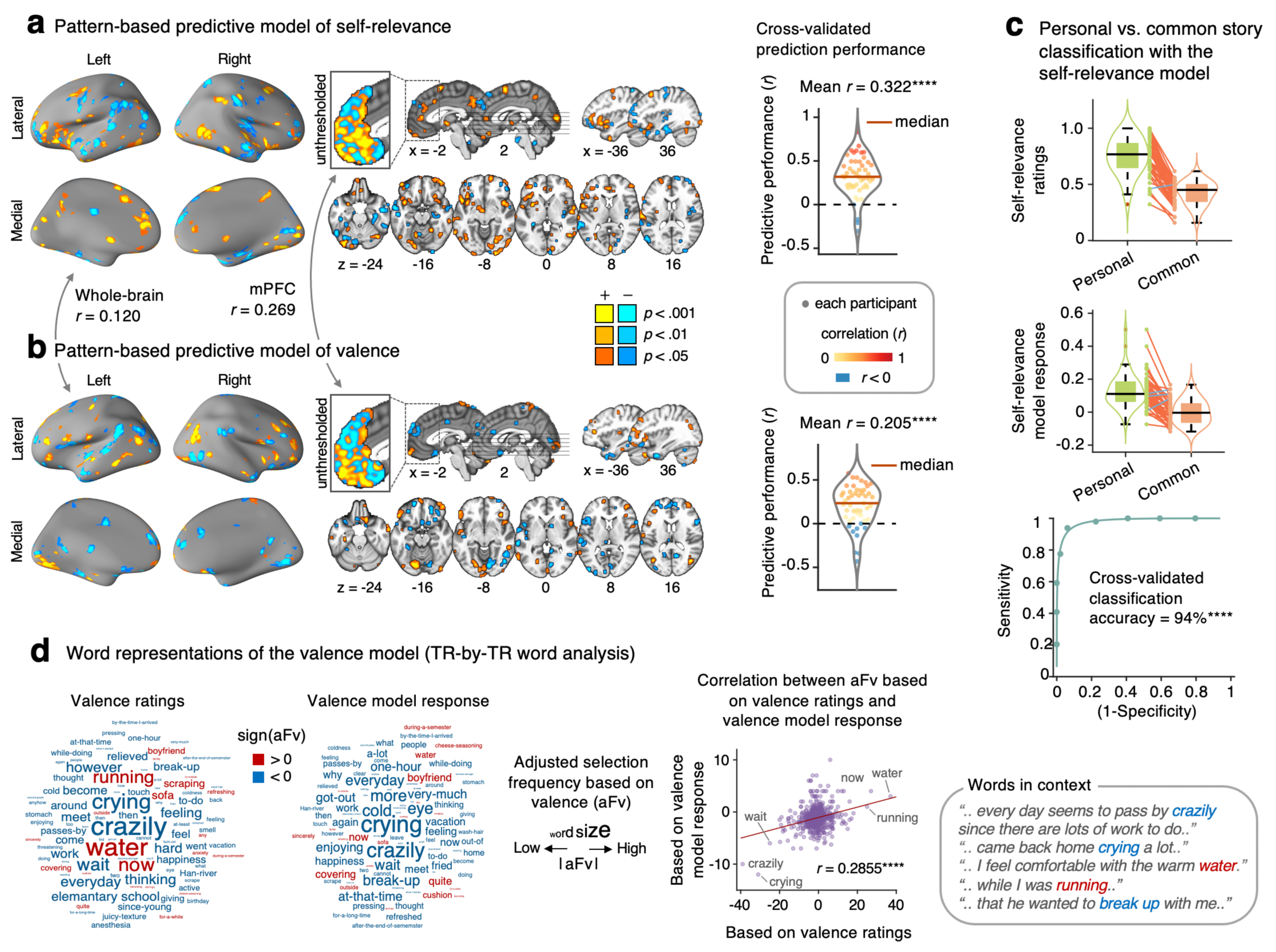
Multivariate pattern-based predictive models of self-relevance and valence. **a,** Multivariate pattern-based predictive model of self-relevance. The brain map shows the predictive weights (positive in warm colors, negative in cool colors) that reliably contributed to the prediction of self-relevance based on bootstrap tests (thresholded at uncorrected *p* < 0.001, two-tailed). We thresholded the map for the purpose of display and interpretation; all weights were used in the prediction. We also pruned the map using two more liberal thresholds, uncorrected *p* < 0.01 and *p* < 0.05, two-tailed, to show the extent of activation clusters. The inset shows the unthresholded weight patterns within the medial prefrontal cortex (mPFC). The violin plot on the right shows the leave-one-subject-out cross-validated model performance, *n* = 49, mean *r* = 0.322, *p* < 2.220e-16, bootstrap test, two-tailed. Each dot indicates the predictive performance of each participant, and the thick red line indicates the median. \*\*\*\**p* < 0.0001. **b,** Multivariate pattern-based predictive models of valence. **c,** (top) The personal stories showed a higher level of self-relevance ratings compared to the common stories, *t*48 = 15.86, *p* < 2.2204e-16, two-tailed, paired *t*-test. orange line: higher scores in the personal stories, blue line: higher scores in the common stories. (middle and bottom) The personal stories showed a higher level of the self-relevance model response than the common stories, *t*48 = 10.18, *p* < 2.2204e-16, and the forced-choice classification accuracy was 93.8%, *p* = 6.980e-11, two-tailed, binomial test. **d,** Word representations of the valence model in TR-by-TR word analysis. We used the adjusted selection frequency based on valence (aFV; see **Methods**) to examine the relative importance of words based on valence ratings or valence model responses. The wordclouds show the relative aFv scores for the top 100 words selected based on the absolute values of actual valence ratings (left) or model responses (middle). The word size represents the absolute magnitude of aFv. The scatter plot shows the relationship between the aFv scores based on the valence ratings (x-axis) and the valence model responses (y-axis) from the whole word set. The ‘words-in-context’ box shows the context in which some top words were used.

**Figure 3.**
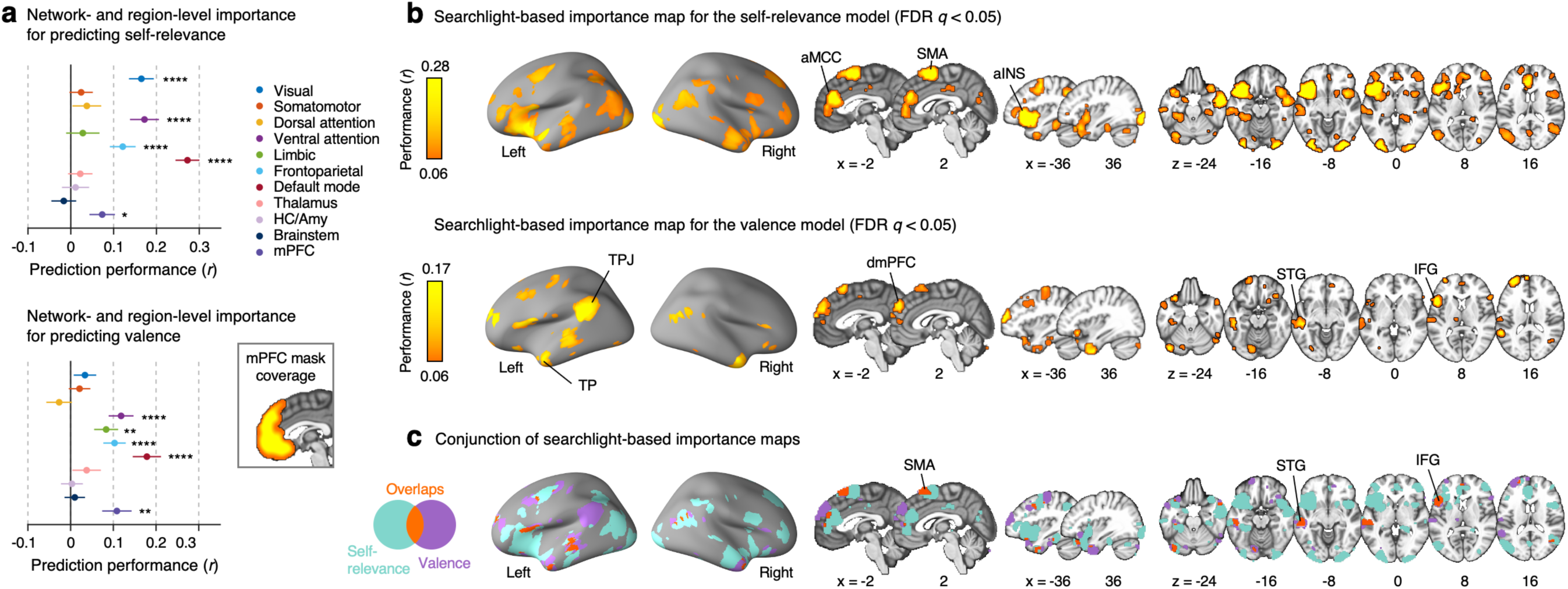
Important features of the self-relevance and valence models (virtual isolation analysis). We examined which features were important for predicting self-relevance and valence using the virtual isolation analysis, which calculated prediction performance (i.e., prediction-outcome correlation) based on a *single* large-scale network, region, or searchlight at a time. **a,** The virtual isolation analysis results for the self-relevance model (top) and the valence model (bottom) with the large-scale networks and some ROIs. Each colored dot represents the prediction-outcome correlations for each network or region with bootstrap tests with 10,000 iterations. The error bars represent the standard deviation of the sampling distribution. \**p* < 0.05, \*\**p* < 0.01, \*\*\**p* < 0.001, \*\*\*\**p* < 0.0001. **b,** Searchlight-based virtual isolation analysis results for the self-relevance model (top) and the valence model (bottom). The maps were thresholded at FDR *q* < 0.05, two-tailed, bootstrap tests. **c,** Conjunction of the searchlight-based importance maps for the self-relevance and valence models. aINS, anterior insula; aMCC, anterior mid-cingulate cortex; dmPFC, dorsomedial prefrontal cortex; IFG, inferior frontal gyrus; SMA, supplementary motor area; STG, superior temporal gyrus; TP, temporal pole; TPJ, temporoparietal junction. For the brain maps of the large-scale networks and ROIs, please see **Supplementary** Fig. 6.

As shown in **Figs. 2a** and **2b**, the two models showed a weak correlation between the unthresholded whole-brain patterns of predictive weights, *r* = 0.120, while the medial prefrontal cortex (mPFC), which is known to be important both for self-referential and valence information processing^40–43^, showed a weak, but larger correlation (*r* = 0.269) than the whole brain, suggesting the existence of overlapping representations between self-relevance and valence within the mPFC. For example, both self-relevance and valence models thresholded at *p* < 0.001 (two-tailed, bootstrap test) showed negative weights within the dorsomedial prefrontal cortex (dmPFC, Brodmann area [BA] 9) and the subgenual anterior cingulate cortex (sgACC, BA25) and positive weights in the ventromedial prefrontal cortex (vmPFC, BA11). However, given that these are the uncorrected results, we conducted further analyses on the network-level and voxel-level importance, the results of which we describe in the next section.

We also conducted additional analyses on the self-relevance and valence models to further examine their validity. First, we tested whether the self-relevance model could classify the personal versus common stories. As **Fig. 2c** shows, the self-relevance ratings were significantly higher in the personal stories (0.75 ± 0.16 [mean ± SD]) than in the common stories (0.42 ± 0.10), *t*48 = 15.86, *p* < 2.220e-16, two-tailed, paired *t*-test, consistent with our assumption that the self-generated personal stories would be more self-relevant than others-generated common stories. The cross-validated responses of the self-relevance model were also significantly higher for the personal stories than the common stories, *t*48 = 10.18, *p* < 2.220e-16, two-tailed, paired *t*-test. The forced-choice classification accuracy with LOSO-CV was 93.8%, *p* = 6.980e-11, two-tailed, binomial test. These results suggest that reading personal and common stories induces different brain representations, and the differences can be captured by our self-relevance model.

Second, to examine the validity of the valence model, we selected the top 20 positive and negative words from the six common story sets based either on the TR-by-TR actual valence ratings or the TR-by-TR valence model responses. We then compared the selection frequency between the ratings versus model responses after adjusting the frequency value with the overall word frequency, which we named adjusted selection frequency based on valence (aFv; for details, see **Methods**). As shown in **Fig. 2d**, the aFv values based on the actual valence ratings showed a significant correlation with the aFv values based on the valence model, *r* = 0.2855, *p* = 1.110e-16. For example, the words ‘crazily,’ ‘crying,’ and ‘wait’ had low aFv values based on ratings and model response, whereas ‘water,’ ‘now,’ and ‘running’ showed high aFv values in both measures. **Fig. 2d** also provides the contexts in which these words were used.

We also conducted univariate general linear model (GLM) analyses for the comparisons (**Supplementary Fig. 5**). Although the univariate maps showed activation patterns distinct from the predictive models (whole-brain pattern similarity between the GLM and relevant predictive model maps (**Supplementary Figs. 5a-b,** *r* = 0.302 for valence and *r* = 0.226 for self-relevance), there were also some consistent findings. For example, the contrast map for the personal vs. common stories (**Supplementary Fig. 5c)** and the parametric modulation map of self-relevance ratings showed strong activations within the anterior midcingulate cortex (aMCC) and anterior insula (aINS), which were consistent with the multivariate pattern-based predictive model of self-relevance (*r* = 0.142 between the contrast map and self-relevance predictive model). In addition, the parametric modulation map of valence ratings showed positive vmPFC activation, consistent with the multivariate pattern-based valence model.

### Network-and region-level importance of predictive models

To achieve the second research goal, i.e., comparing brain representations of self-relevance and valence (**Fig. 1a**), we further examined the importance of various features for the self-relevance and valence models^44^. Specifically, we examined network-and region-level importance with virtual isolation analysis (i.e., calculating the prediction-outcome correlation based on a *single* large-scale network, region, or searchlight at a time) the virtual lesion analysis^44^ (i.e., calculating the changes in the prediction-outcome correlation after *removing* one network, region, or searchlight at a time). As shown in **Fig. 3a**, the virtual isolation analysis using large-scale networks and some regions-of-interest (ROIs) showed that the default mode, ventral attention, and frontoparietal networks had significant prediction performances for both self-relevance and valence (bootstrap test with 10,000 iterations; for details, see **Supplementary Table 1**). When we tested the mPFC region separately, it also showed significant prediction for both self-relevance and valence. The visual network was, however, important only for predicting self-relevance, while the limbic network was important only for predicting valence. The virtual lesion analysis using large-scale networks and ROIs also showed that the default mode network (DMN) was important for predicting both self-relevance and valence (see **Supplementary Fig. 7a** and **Supplementary Table 1**), while the visual, ventral attention, and frontoparietal networks were important only for predicting self-relevance, and the mPFC was important only for predicting valence. For the brain maps of the large-scale networks and ROIs, please see **Supplementary Fig. 6**.

**Fig. 3b** and **Supplementary Fig. 7b** present the searchlight analysis results for the virtual isolation and virtual lesion analyses, respectively. The searchlight-based virtual isolation results show that the aMCC, aINS, and visual areas were important for the prediction of self-relevance, and for the valence model, the dmPFC, temporo-parietal junction (TPJ), temporal pole (TP) and other regions were important predictors. **Fig. 3c** shows the conjunction map between the importance maps of self-relevance and valence models, and the supplementary motor area (SMA), superior temporal gyrus (STG), inferior frontal gyrus (IFG) appeared to be important for both models.

### Decoding the levels of self-relevance and valence during free-thinking and resting

To achieve the third research goal, i.e., decoding the levels of self-relevance and valence during free-thinking and resting (**Fig. 1a**), we first tested the models on the fMRI data from the thought-sampling task, in which we asked participants to report what they were thinking every 50 seconds with jitters (interval = 50.7 ± 5.6 [mean ± SD] seconds; see **Methods** for the details of the task). Given that the verbal responses to thought sampling are most likely to be based on the thought contents just before the reporting onset and considering the hemodynamic delay, as shown in **Fig. 4a**, our time-of-interest was a 10-TR (a total of 4.6 seconds) time window around the reporting onset. We evaluated the prediction performance with the prediction-outcome correlations with LOSO-CV using the moving-window approach based on the data convolved with the temporal Gaussian kernel (FWHM = 10 TRs). As in **Fig. 4b** left panel, both predictions of self-relevance and valence showed the best prediction performance at the time-of-interest period. When we examined the prediction performances of time bins of 10 TRs (**Fig. 4b** right panel), both models showed weak but significant correlations for the corresponding ratings at the time-of-interest bin (for the self-relevance model predicting self-relevance ratings, mean *r* = 0.0518, *p* = 0.0141, one-tailed, bootstrap test with 10,000 iterations; for the valence model predicting valence ratings, mean *r* = 0.0495, *p* = 0.0095). To compare our model performance with other pre-defined models, we also tested 9 *a priori* maps from previous studies in the same manner. The *a priori* maps included two component maps of self-generated thought^45^, six meta-analytic maps of aversion, episodic, default, self, emotion, and semantic from referenced study^46^, and the picture-induced negative emotion signature (PINES)^47^ (**Fig. 4c**). The result showed that only the models from this study were predictive of the self-relevance and valence ratings. We also examined which large-scale networks and ROIs were important for these predictions with the virtual isolation analysis and found that the DMN was the predictor important for both self-relevance and valence models (**Supplementary Fig. 8**).

**Figure 4.**
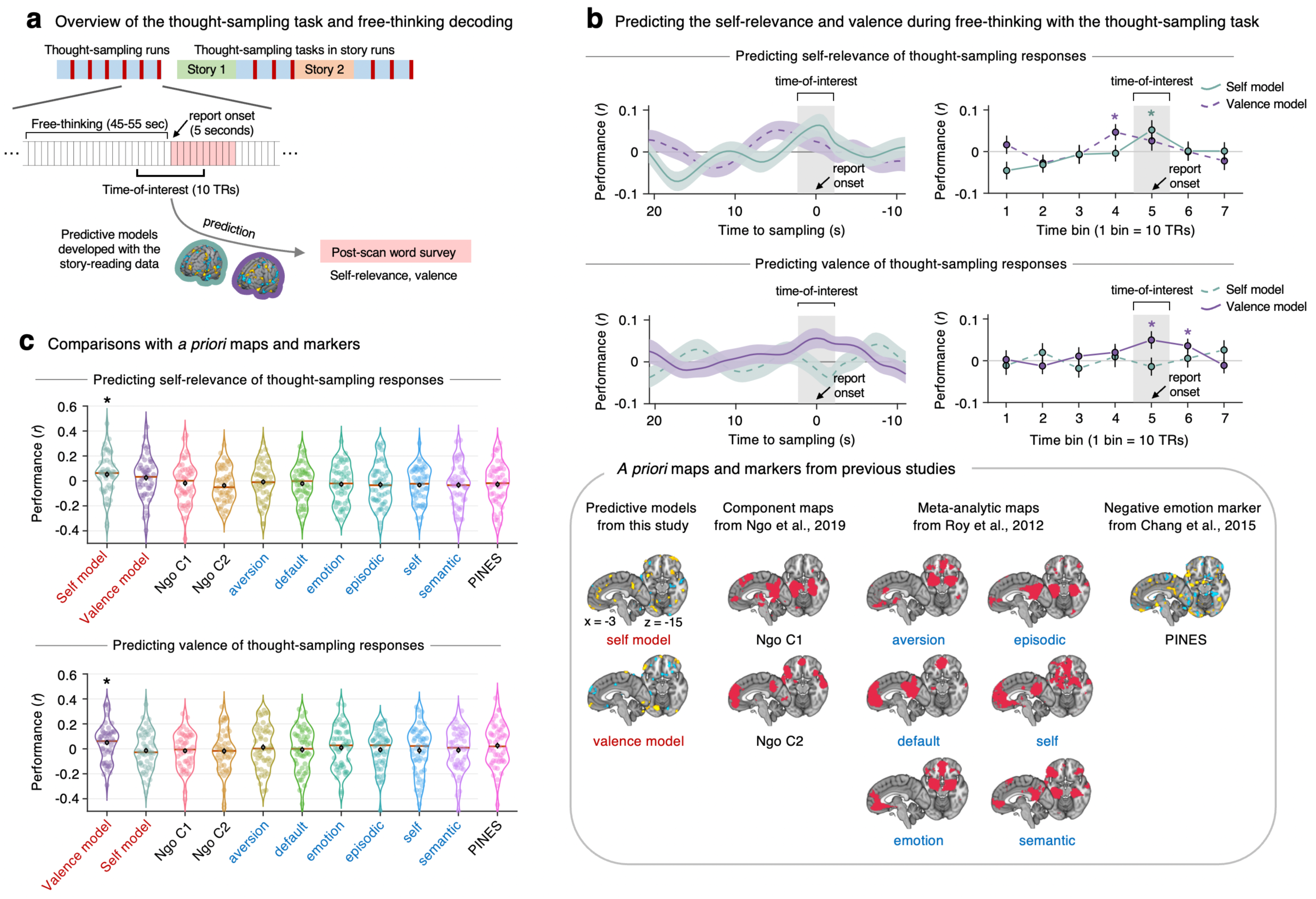
Decoding self-relevance and valence during free-thinking. **a,** During the thought-sampling task, participants were instructed to think freely for around 50 seconds and then verbally report what they were thinking in words or phrases. One thought-sampling run included a total of 6 trials, and each story-reading was followed by 3 trials of thought-sampling. We applied the predictive models of self-relevance and valence to the fMRI data from the free-thinking period to predict the self-relevance and valence ratings collected from the post-scan survey. **b,** Model performances (top: predicting self-relevance, bottom: predicting valence) measured by the prediction-outcome correlations. The plots on the left show the prediction performance using a moving-window approach based on the data convolved with a temporal Gaussian kernel (FWHM = 10 TRs). The plots on the right show the prediction performances for the time bins of 10 TRs. Shading and error bars indicate the standard deviation of performances across all participants. **c,** The prediction performances of the self-relevance and valence models from the current study and 9 a priori maps (the a priori maps are shown on the right). The *a priori* maps included 2 component maps of self-generated thought^45^, 6 meta-analytic maps ^46^, and the picture-induced negative emotion signature (PINES)^47^.

Lastly, we tested the self-relevance and valence models on the resting-state fMRI data from an independent dataset (*n* = 90), in which, at the end of the 6-minute resting scan, participants reported the levels of self-relevance and valence for the thoughts they had during the resting scan (post-resting survey; **Fig. 5a**). Given that the answers to the post-resting survey were likely to be based on participants’ thoughts around the end of the scan, we tested the model performances using averaged data from the end of the scan. As shown in **Fig. 5b**, the self-relevance and valence models showed significant predictions for the self-relevance and valence ratings around the same temporal window, which was the 27^th^ to 33^rd^ TRs (for self-relevance) and the 26^th^ to 37^th^ TRs (for valence) from the end of the scan (the gray boxes in **Fig. 5b**, which indicated the time points with *r* > 0 and *p* < 0.05, one-tailed, one-sample *t*-test). The best prediction was observed when the data of the last 31 TRs (i.e., 14.3 seconds) were averaged, and the prediction performance were *r* = 0.189, *p* = 0.037 for the self-relevance model and *r* = 0.300, *p* = 0.002 for the valence model (one-tailed, scatter plots in **Fig. 5b**). Similar to what we did for the free-thinking decoding, we examined which large-scale networks and ROIs were important for these predictions with the virtual isolation analysis. For self-relevance, only the DMN was significant, while for valence, the ventral attention and limbic networks, and brainstem were significant. (**Fig. 5c**)

**Figure 5.**
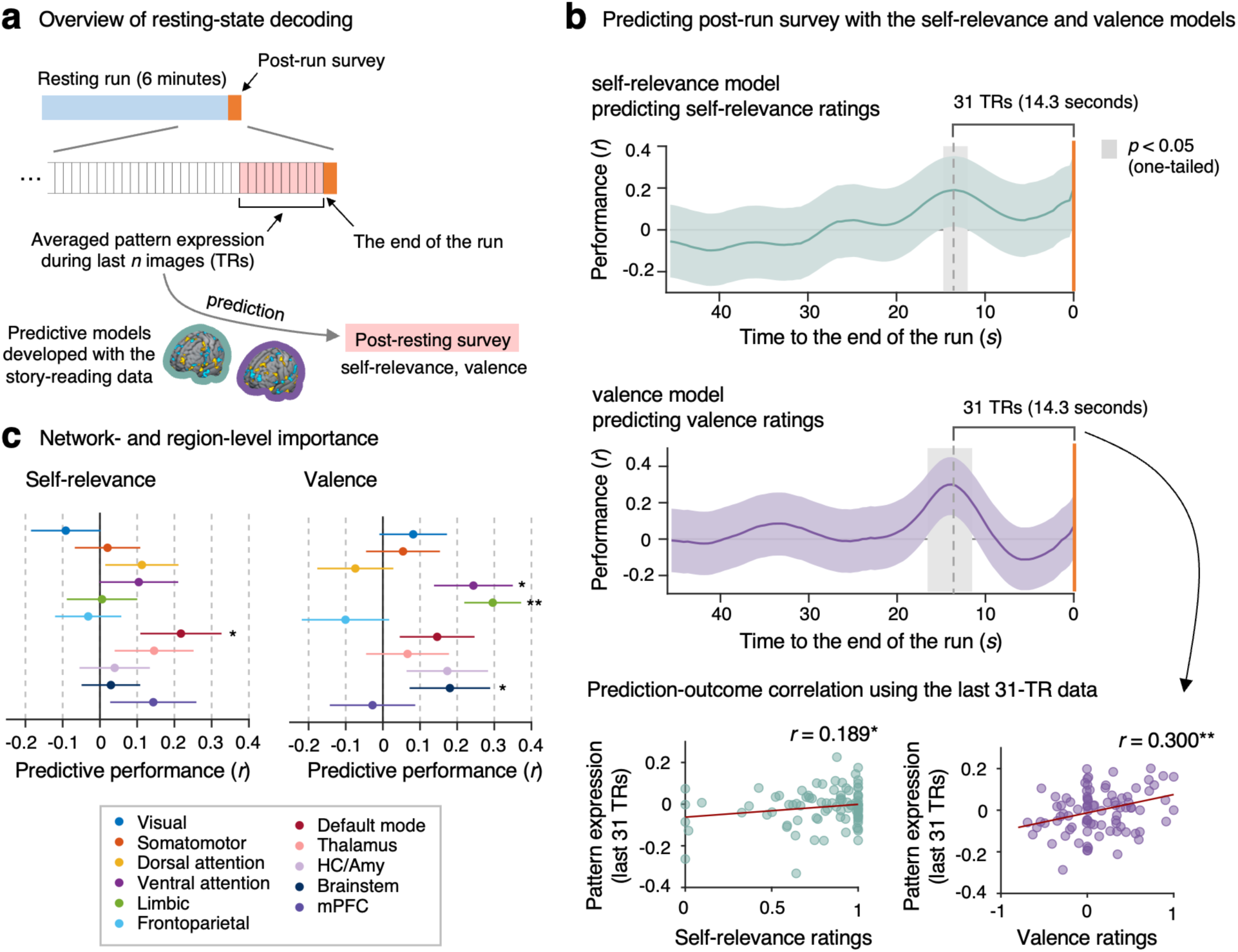
Decoding self-relevance and valence during rest. **a,** We tested our models on data from a six-minute resting-state run from an independent dataset (*n* = 90) to predict self-relevance and valence ratings from a post-run survey on spontaneous thoughts during resting. **b,** Prediction performance was calculated as the prediction-outcome correlation based on the averaged data from the end of the scan. The gray boxes indicate the time period where each model showed significant positive predictions (i.e., *p* < 0.05 and *r* > 0). The scatter plots show the relationship between the ratings and pattern expression values when the last 31 TRs (14.3 seconds) were averaged. Each dot indicates each participant. \**p* < .05, \*\**p* < 0.01, one-tailed, one-sample *t*-test. **c,** Network-and region-level decoding performance. Each colored dot represents the mean prediction-outcome correlations for each network or region with bootstrap tests with 10,000 iterations. The error bars represent the standard deviation of the sampling distribution, \**p* < .05, \*\**p* < 0.01, one-tailed, bootstrap test.

## Discussion

In this study, we developed multivariate pattern-based predictive models of self-relevance and valence that can be used to decode spontaneous thoughts. For this, we conducted an fMRI experiment using the story-reading task, in which we showed personal stories created through one-on-one online interviews with participants and common stories created by others. The newly developed self-relevance and valence models could decode the levels of these two content dimensions of spontaneous thought during rest across two datasets. The main innovations and findings of this study can be summarized as the following: (1) we used personal stories as stimuli to evoke thoughts and emotions that resemble spontaneous thoughts; (2) we identified brain systems important for decoding self-relevance and valence using virtual isolation and lesion analyses, such as the mPFC, aINS, and TPJ; and (3) we were able to decode the level of self-relevance and valence during rest with or without thought sampling using the two predictive models.

First, we used personal narratives as experimental stimuli to induce cognitive and emotional states similar to spontaneous thoughts. We used self-generated personal stories as stimuli for the following two reasons: (1) self-relevant thoughts, such as current personal concerns, past memories, and future plans, are known to be the major building blocks of spontaneous thought^7, 24, 25^, and (2) recent studies suggested that spontaneous thoughts are experienced in the form of deeply processed imagery and concepts, such as narratives^16, 17^. Therefore, personal stories based on participants’ past experiences and related emotions should share many characteristics with spontaneous thoughts. In addition, following recent efforts in the neuroimaging field to use naturalistic stimuli, such as movies^23, 48, 49^, we created individually unique stories through one-on-one interviews with participants to make the experimental stimuli as natural as possible. The common stories used for all participants were also created by pilot participants through the same one-on-one interview procedure. Therefore, we believe our story stimuli were able to induce natural and vivid thoughts and emotions that resemble spontaneous thoughts during the scan. Using these stories as a bridge between the experiment and naturally experiencing spontaneous thoughts, we could develop fMRI-based predictive models that could decode not only the contents of current reading materials but also the contents of individuals’ spontaneous thoughts.

Second, we identified brain regions and networks important for decoding the levels of self-relevance and valence. Using the virtual isolation and lesion methods^44^, we found some converging evidence that the DMN, including the mPFC, plays an important role in predicting both self-relevance and valence. In particular, for decoding levels of self-relevance and valence during free-thinking, the DMN was the only significant predictor both for self-relevance and valence (**Supplementary Fig. 8**). This is consistent with previous literature suggesting that the DMN is important for processing details of ongoing cognition^15^, memory^50, 51^, and self^52, 53^. There were also brain networks and regions that were uniquely important for each model. For example, the visual network (**Fig. 3a**), aINS, and aMCC (**Fig. 3b**) were among the important features for decoding self-relevance, but these were not important for valence. Conversely, the limbic network (**Fig. 3a**) and the left TPJ and dmPFC (**Fig. 3b**) were among the important features for decoding valence, but these were not important for self-relevance. The importance of the aINS and aMCC for self-relevance can be understood in the context of the salience network (or the ventral attention network), considering that the aINS and aMCC are among the key hubs of the network. The salience network is known to be important for context-dependent salience detection^35, 36^, and it is likely that, when there is no external task (such as reading autobiographical stories without external tasks), the self can become the most important context, which is generated from within. For example, in our task, the primary concern or ongoing implicit task could become identifying content relevant to oneself. This could be the case for many other naturalistic situations where we do not have external tasks. In addition, the importance of the limbic network in emotional valence is also consistent with previous studies reporting that the orbitofrontal cortex (OFC) and temporal pole (TP), which are the main components of the limbic network, were important for valence processing ^43, 54^. We also found that the left TPJ, along with the dmPFC (see **Fig. 3b**), were important for valence. This finding was rather unexpected, but the left TPJ and dmPFC are the major constituents of DMN’s dorsal medial subsystem^55^. This subsystem has been suggested to play an important role in a high-level “mind’s mind” form of imagination, such as reflective thinking in the verbal form^56^. Thus, we can speculate that feeling positive or negative emotions from reading autobiographical stories requires more mind’s mind form of imagination, such as mentalizing (cf. ref. ^57^).

Third, with our self-relevance and valence models, we were able to decode the respective content dimension scores during free-thinking and resting. The models showed weak but significant prediction performances across two different datasets with two different task contexts—i.e., thought sampling and resting. Given that the contents and dynamics of spontaneous thought could provide rich information about individuals’ mental and brain health, the ability to decode some aspects of spontaneous thought directly from neuroimaging data should be useful. This represents a powerful combination of task-based data (i.e., personal and common stories) with annotated rest in an effort to decode the contents of the mind during rest using a known ground truth^58^. In this study, we focused on two content dimensions of spontaneous thought—self-relevance and valence, which are among the important predictors of depression and negative affectivity scores in sub-clinical populations^6, 7^. Though further studies, including clinical ones, are needed to identify which content dimensions are most useful for predicting and promoting mental health and well-being, this study showed the potential to use fMRI data to extract certain information about spontaneous thought. Importantly, we demonstrated that resting-state fMRI could be used to decode aspects of spontaneous thought, opening a new avenue for using resting-state data involving no or minimal tasks (e.g., introspection) to obtain rich information about one’s internal cognition. This is important because it is often difficult (or impossible) to administer task-based fMRI tests to patients. In addition, our study further implies that we should reconceptualize the notion of the resting state. As we wrote in the introduction, our mind (and brain) never rests. Our mind keeps wandering, and our brain keeps being activated spontaneously. Therefore, the resting state should be reconceptualized as spontaneous cognition or spontaneous activity conditions. With this new perspective, we will be able to develop new tasks that are based on resting but have minimal task components, which then can be used to provide richer and clinically more useful information.

There are some limitations in this study. First, we used two different types of stories— personal and common stories to increase the variance of the level of self-relevance, the orthogonality between self-relevance and valence, and the comparability across participants. However, it is also possible that they are qualitatively different and thus might not be comparable between the two conditions. We made multiple efforts to resolve this issue. Given that we expected personal stories to be more familiar than common stories, we had participants read all common stories on Day 1. We also gave quizzes on the common stories twice—first, on Day 1 after reading the stories, and second, on Day 2 right before the fMRI experiment. In addition, we collected the ratings of concentration and familiarity for each story on a scale of 0 to 1 after each run. The results showed that the concentration and familiarity ratings for the personal stories (0.78 ± 0.17 [mean ± SD] and 0.87 ± 0.16, respectively) were still higher than the common stories (0.69 ± 0.21 and 0.74 ± 0.19), indicating that personal stories elicited higher levels of concentration and familiarity. Nevertheless, the concentration and familiarity ratings for common stories were also high enough for us to assume that the two types of stories were not qualitatively different. In future studies, however, it would be good to try using a single type of story with variable levels of self-relevance and valence. Second, the prediction performances for the decoding of free-thinking and resting-state data were quite low. This may suggest that the brain representations of spontaneous thought are highly complex and idiosyncratic. In a recent study, we showed that the brain representations of valence become increasingly idiosyncratic as thought topics become more self-relevant^6^. Therefore, adopting a personalized modeling approach to studying spontaneous thought (e.g., extensive sampling of small*-*N design) could be considered in future studies. Third, the current findings need further replications. Though we showed generalizable predictions across two independent datasets, the prediction performances were not high. Thus, more replications will be needed to support the generalizability of our findings.

Overall, this study used a novel approach to the brain decoding of spontaneous thought—using self-generated personal stories to develop brain-based predictive models of self-relevance and valence of spontaneous thought contents. As these models showed the potential to decode these spontaneous thought content dimensions during free-thinking or resting state, this study provides an important step toward developing brain models of internal thoughts and emotions during daydreaming.

## Author Contributions

H.J.K. and C.-W.W. conceived, designed, and conducted the experiment. B.K.L. provided the independent test dataset for resting-state decoding. H.J.K. and C.-W.W. analyzed the data, interpreted the results, and wrote the manuscript. E.S.F. and C.-W.W. edited the manuscript and provided supervision.

## Acknowledgments

This work was supported by IBS-R015-D1 (Institute for Basic Science; to C.-W.W.) and 2021M3E5D2A01022515 (National Research Foundation of Korea; CWW).

## Data Availability

The data used to generate figures, including the predictive models, will be shared upon publication through a Figshare repository.

## Code Availability

The codes for generating the main figures will be shared upon publication through a Figshare repository. In-house Matlab codes for fMRI data analyses are available at https://github.com/canlab/CanlabCore and https://github.com/cocoanlab/cocoanCORE.

## Methods

### Participants

Fifty-seven healthy right-handed participants completed the experiment. Participants provided written informed consent in compliance with the guidelines of the Sungkyunkwan University Bioethics Committee and the approval from the institutional review board (IRB). Participants with psychiatric, neurological, or systemic disorders and MRI contraindications were excluded. All participants had a normal or corrected-to-normal vision and were naïve to the purpose of the experiment. All participants were Koreans and spoke Korean as their first language. All experimental procedures were conducted using the Korean language. We included forty-nine participants (age = 22.8 ± 2.4 [mean ± SD], 21 female) in the final data given that we excluded 8 participants total (6 participants due to poor performance [e.g., did not focus on the task, sleep during the scan, or did not fully understand the task], and 2 participants due to poor image quality).

### Experimental procedure

The experiment consisted of three components across two-day sessions (**Fig. 1b**): (1) an online interview on Day 1, (2) an fMRI experiment with the story-reading and thought-sampling tasks on Day 2, and (3) a post-scan survey on Day 2. First, on Day 1, after providing an overview of the experiment and having participants complete a pre-scan survey of self-report questionnaires, we conducted a one-on-one interview to create personal stories (for the details of the interview and story-making procedure, please see the “Personal and common story sets” section below and **Supplementary Fig. 9**). After the interview, the participants read the common stories to match the level of familiarity between the personal and common story sets. Then, participants visited the lab again about one week (7.32 ± 2.8 [mean ± SD] days) after their first visit for the fMRI experiment and post-scan survey. The fMRI experiment had a total of 7 runs, which consisted of 5 story-reading runs (about 14 minutes per run) and 2 thought-sampling runs (about 6 minutes per run). We placed the thought-sampling runs in the first and last runs of the 7 runs. We used MATLAB (Mathworks) and Psychtoolbox (version 3.0.16, http://psychtoolbox.org/) for stimuli presentation and behavioral data acquisition. After the fMRI experiment, we conducted a post-scan survey (see the “Post-scan survey” section below).

### Personal and common story sets

To create personal stories, we conducted a one-on-one interview with each participant on Day 1. The interview was conducted in the laboratory, but the interview was performed using a computer through an online messenger to help participants feel more comfortable about sharing their personal stories. We used the KakaoTalk (Kakao Inc.) platform, which is currently the most popular messenger application in South Korea, and young Korean people are used to sharing their thoughts and feelings through the application. During the interview, we asked participants to share past memories and personal feelings related to the following four topics: 1) pain and suffering, 2) pleasure and joy, 3) danger and threat, and 4) safety and comfort. This resulted in four unique sets of story ingredients for each participant. We described the details of the procedure of the interview in **Supplementary Fig. 9**. After the interview, we combined the story ingredients to make story stimuli have a good length (202 ± 6 [mean ± SD] characters, which corresponds to 222.8 ± 1.5 seconds of the screen display) and to make them better suited for screen display (e.g., correcting typos and removing blanks, etc.). Then the participants re-read the modified version of the stories and confirmed that the modification did not change the content. In addition to the four personal stories, we made a common set of story stimuli that were the same across all participants (**Supplementary Fig. 10**). We created the common stories with participants of the pilot study and used the same four topics described above, but we also had two additional topics: (5) relationships and love and (6) break-ups and hatred. The length of the common stories was 193 ± 4 characters and 223.8 ± 0.7 seconds of the screen display. Participants also read the common story set after the interview to become familiar with the common stories. This step is important to keep the levels of familiarity similar between the personal and common stories. To ensure that participants became familiar with the common story contents, we administered quizzes on the common stories twice—first, on Day 1 after the interview, and second, on Day 2 right before the fMRI experiment. On Day 1, after participants read the common stories, we asked two true-false questions for each common story, and the average score was 11.65 ± 0.60 [mean ± SD] out of 12 questions, suggesting that the participants understood the common stories well. In addition, before the fMRI experiment on Day 2, we asked one question for each common story, and the average score was 4.98 ± 0.95 out of 6, which shows that participants remembered the common stories well.

### Story-reading task

In the story-reading task, participants read a total of 10 stories, which consisted of four personal stories and six common stories. We included two stories in one run and made those two stories to be heterogeneous in terms of story types (i.e., personal versus common) and the emotional valence of topics to include a wider distribution of our modeling targets in our training data. To this end, in four runs, we presented one personal and one common story, and in one remaining run, we presented two common stories. The orders of stories and runs were counterbalanced across participants. In addition, in each run, we presented one story from positively valenced topics (e.g., ‘pleasure and joy,’ ‘safety and comfort,’ ‘relationships and love’) and one story from negatively valenced topics (e.g., ‘pain and suffering,’ ‘danger and threat,’ ‘break-ups and hatred’). For the story-reading task, we showed one word at a time at the center of the screen based on a rapid serial visual presentation (RSVP) procedure^59^ (see **Fig. 1b**). Each word was displayed on the screen for a jittered time with 0.96 ± 0.20 (mean ± SD) seconds. There was an additional pause after a comma (1.0 seconds) or a period (1.5 seconds) to provide enough time to process the story. Each story included 201.92 ± 6.10 words on average, and the average time of story presentation was 222.80 ± 1.45 seconds. During the story-reading, we asked three valence questions (i.e., current emotional valence) intermittently. After each story-reading, we had a free-thinking period with the thought-sampling task for 150 seconds, which included three trials of thought sampling (for details of the thought-sampling task, see the next section).

### Thought-sampling task

In the thought-sampling task, each trial consisted of a free-thinking period, which lasted for around 50 seconds (50.7 ± 5.6 seconds), and a thought-sampling period, which was around 5 seconds. One thought-sampling run included a total of 6 trials, and each story-reading was followed by 3 trials of thought-sampling. During the free-thinking period, participants were instructed to think freely while looking at the fixation point displayed on the screen. After the free-thinking period, thought sampling started with a question, “Please report what you are thinking now in words or phrases,” which appeared on the screen for 5 seconds (**Fig. 1b**). The participants then verbally reported words or phrases that captured the essence of their current thought contents. Participants’ verbal reports were recorded and transcribed by the experimenters through an MR-compatible microphone with a noise-cancellation function (FOMRI, Optoacoustics Inc.). After each run, we showed the transcribed responses to the participants and obtained their confirmation of whether the transcription was correct. A total of 42 responses were collected per participant—6 responses from two thought-sampling runs and 3 responses after 10 story-reading. These reported responses were used in the post-scan word survey.

### Post-scan survey

After the fMRI scan, participants performed a post-scan survey for thought-sampling responses (i.e., words or phrases) and stories in the behavioral experimental room (**Fig. 1c**). For each thought-sampling response, we asked participants to rate the response on five content dimensions, including (1) valence (whether the response was negative or positive), (2) self-relevance (how much the response was related to themselves), (3) perceived time (which time point was related to the response), (4) vividness (how vivid was the response), (5) safety/threat (whether the response was perceived as safe or threatening). For the story survey, we asked participants to rate the stories on three content dimensions, including valence, self-relevance, and vividness. Because vividness was not the primary focus of this study, it was collected from only four out of ten stories, and thus, we did not analyze the vividness ratings in the current study.

Sentences of the story were presented on the screen with a horizontal line below them. Participants used a tablet pen to draw a continuous curve that provided their ratings about each part of the stories.

### fMRI data acquisition and preprocessing

Whole-brain MRI data were acquired on a 3T Siemens Prisma scanner at Sungkyunkwan University with a 64-channel head coil. High-resolution T1-weighted structural images were acquired with TR = 2400 ms and TE = 2.34 ms. Functional EPI images were acquired with TR = 460 ms, TE = 27.2 ms, multiband acceleration factor = 8, field of view = 220 mm, 82 × 82 matrix, 2.7 × 2.7 × 2.7 mm^3^ voxels, 56 interleaved slices. The number of volumes was 812 for the thought-sampling runs and 1,855 for the story-reading runs. All preprocessing steps were performed using SPM12, FSL, and MATLAB. Before preprocessing, 18 initial volumes of fMRI data were removed to allow enough time for image intensity stabilization. Structural T1-weighted images were co-registered to the functional image and normalized to the MNI standard brain space. Functional EPI images were motion-corrected (realigned), distortion-corrected, and normalized to MNI using T1 images with the interpolation to 2 × 2 × 2 mm^3^ voxels. Then we smoothed the functional images with a 5 mm FWHM Gaussian kernel. To identify spike-related noise in fMRI data, the Mahalanobis distance of each time point from every voxel in each run was calculated. Time points with Mahalanobis distance above 3 standard deviations from the mean were considered spikes. Twenty-four head motion parameters (6 movement parameters including x, y, z, roll, pitch, and yaw, and their mean-centered squares, derivatives, and squared derivatives) were estimated, and all images were realigned to the first functional image using interpolation according to motion data. The nuisance regressors consisted of *n* regressors from spike identification (the number of regressors depends on the number of spikes), 24 regressors from motion data, 1 for linear drift, 5 top principal components from white matter signal, and 5 top principal components from CSF signal. Also, the timings that would involve task-related motions (e.g., speaking moments in the thought-sampling task, valence ratings during story-reading, and post-story ratings during scanning) were also included as an additional regressor to filter out task-dependent artifacts. For the thought-sampling runs, which were similar to resting-state runs, we used a bandpass filter (0.008 Hz -0.1 Hz) to reduce physiological noise and conducted nuisance regression with run-specific regressors. Also, we winsorized the values outside of the median ± 5 standard deviations (across all the voxels and time points). For the story-reading runs, which are event-related fMRI, we applied a high pass filter with a frequency of 0.001 Hz and skipped the winsorizing step given that it can remove some high-frequency task-related brain signals.

### General linear modeling (GLM) analysis

We conducted the voxel-wise general linear modeling (GLM) analysis with parametric modulation for the story-reading fMRI data. First, the boxcar regressors, convolved with the canonical hemodynamic response function, were constructed to model the word display. We then included three additional regressors that are parametrically modulated with the number of characters in each word, the level of self-relevance, and the level of valence. With these regressors, we conducted a multiple regression. The blank period was used as an implicit baseline. With the beta maps from the first-level GLM analysis, we conducted the second-level analysis with robust regression.

### Data quantization for predictive modeling

To effectively model self-relevance and valence, which are known to be correlated, we concatenated and quantized the data based on quintiles of the two dimensions, constructing 25 averaged images per participant (i.e., 5 levels of self-relevance × 5 levels of valence; **Supplementary Fig. 2**). In more detail, in each participant, we first concatenated fMRI data of five story-reading runs. In this procedure, we only used TRs from story-reading tasks—in other words, before the data concatenation, we removed TRs from the rating and thought-sampling task periods. Considering the hemodynamic delay, all the fMRI data were delayed by 10 TRs (= 4.6 seconds). For each TR of fMRI data, we assigned the self-relevance and valence ratings from the post-scan story survey based on the display onset timing of each word from the stories. We then divided the TR-level images into quintiles based on ratings, resulting in each TR being assigned to 5 levels of self-relevance and 5 levels of valence. Combining data from the same levels of self-relevance and valence, we created 25 (5 levels of self-relevance ratings × 5 levels of valence ratings) possible segments of fMRI and rating data for each participant. We then averaged fMRI and rating data for each segment and used them for predictive modeling.

### Predictive modeling

For predictive modeling, we used the 25 data segments from 49 participants’ data, resulting in a total of 1,225 data points. However, depending on the rating distributions, some segments had no data, resulting in a total of 1208 data segments. We conducted principal component regression (PCR) for training multivariate pattern-based predictive models, in which the fMRI data were the independent variables, and the ratings were the dependent variable. We did not conduct further hyperparameter tuning (e.g., reducing the number of components, etc.) to minimize potential biases in the results. Thus, we used the full component (i.e., 1,208 components) for modeling. To validate the performance of each model, we used two types of cross-validation methods: leave-one-subject-out cross-validated (LOSO-CV) and random-split cross-validation (RS-CV^38, 39^). For LOSO-CV, we obtained a predictive model using data that excluded one participant (i.e., 5 × 5 data points) from the total sample, then used the data from the hold-out participant to test the model. To test whether the observed model performance (i.e., prediction-outcome correlations) was significantly greater than zero, we conducted bootstrap tests on the within-participant prediction-outcome correlations with 10,000 iterations. We also conducted permutation tests to examine whether the observed model performance was greater than its null distribution. For this, we first shuffled the ratings within each participant. We then trained the models with the shuffled data after excluding one participant’s data and tested the models on the held-out participant’s non-shuffled data. We repeated this procedure across all participants (i.e., leave-one-participant-out cross-validation), and the prediction-outcome correlations were averaged across participants for each iteration. We repeated this procedure 1,000 times. The *p*-values were calculated as the probability of obtaining the actual mean prediction-outcome correlation from the null values. For RS-CV, 20% of the participants’ data was randomly selected as the hold-out data for each iteration. Then, we obtained a predictive map using data from the remaining 80% of the participants, and the map was tested to the hold-out data iteratively. This procedure was repeated 50 times.

### Large-scale networks and regions-of-interest (ROIs)

The large-scale functional brain networks included 7 networks within the cerebral cortex^60^, cerebellum^61^, and basal ganglia^62^. The ROIs included the thalamus, hippocampus, and amygdala from the SPM anatomy toolbox^63^ and the brainstem (**Supplementary Fig. 6**).

### Adjusted selection frequency based on valence (aFv)

To investigate the word representations of the valence model, we first selected the top and bottom 20 words from each common story based on the actual or predicted valence ratings and then calculated the selection frequency over the base frequency of the word across common stories. More specifically, we selected the top 20 (most positive) and bottom 20 (most negative) words per participant based on the actual valence ratings in six common stories (a total of the top 120 and bottom 120 words per participant) and then counted the selection frequency for each word in the set. To adjust the base rate of the word frequency, we divided the counts by each word’s base frequency across common stories. We named the value as adjusted selection frequency based on valence (aFv), which is defined as

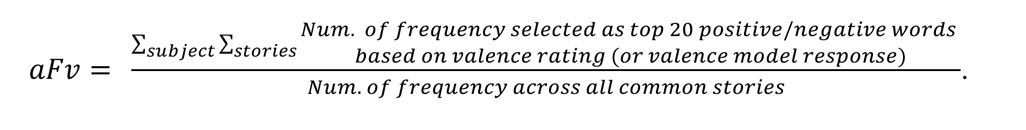

The aFv was calculated based on either actual valence ratings or valence model response (i.e., predicted valence ratings). Based on the aFv, the top 100 words were selected based on the absolute value of actual or predicted valence ratings. **Fig. 2d** shows the aFv distribution of the selected top 100 words based on the actual (left) and predicted (right) valence ratings as wordclouds. The sizes of each word in the wordcloud indicate the levels of aFV, and the color indicates the sign of valence (i.e., positive or negative). The correlation between the aFv based on the actual and predicted valence ratings from the whole word set is shown in the scatter plot.

### Searchlight analysis

We conducted a searchlight analysis for the virtual isolation and lesion analyses (**Fig. 3b** and **Supplementary Fig. 7b**). For the searchlight analysis, we constructed spherical searchlights with a radius of 5 voxels (= 10 mm) around center voxels. The average number of voxels included in each searchlight was 435.9 ± 84.5 (mean ± SD). For the virtual isolation analysis, we only used the predictive weights within searchlights to predict self-relevance or valence ratings. For the virtual lesion analysis, we removed the predictive weights within searchlights, predicted self-relevance or valence ratings with the remaining predictive weights, and calculated the changes in the prediction performance. We used LOSO-CV for all searchlight analyses (i.e., using the predictive weights from the model trained after removing one participant’s data and testing the predictive weights on the left-out participant’s data).

### Independent test dataset for the resting-state decoding

The independent test dataset for the resting-state decoding was from a previous study focusing on the free-association-based thought-sampling task^6^. We tested our models on data from one six-minute resting-state run from the study. The previous study conducted an fMRI experiment on 90 right-handed healthy participants (age = 23.0 ± 2.4 [mean ± SD], 44 female) at the same imaging center as in the current study. The data acquisition parameters and preprocessing pipeline of this independent test dataset were the same as in the current study.

