## Supplementary information for "Getting Personal: Brain Decoding of Spontaneous Thought Using Personal Narratives"

**Running Head:** BRAIN DECODING OF SPONTANEOUS THOUGHT

**This PDF file includes:**

1. Supplementary Figures 1-10
2. Supplementary Table

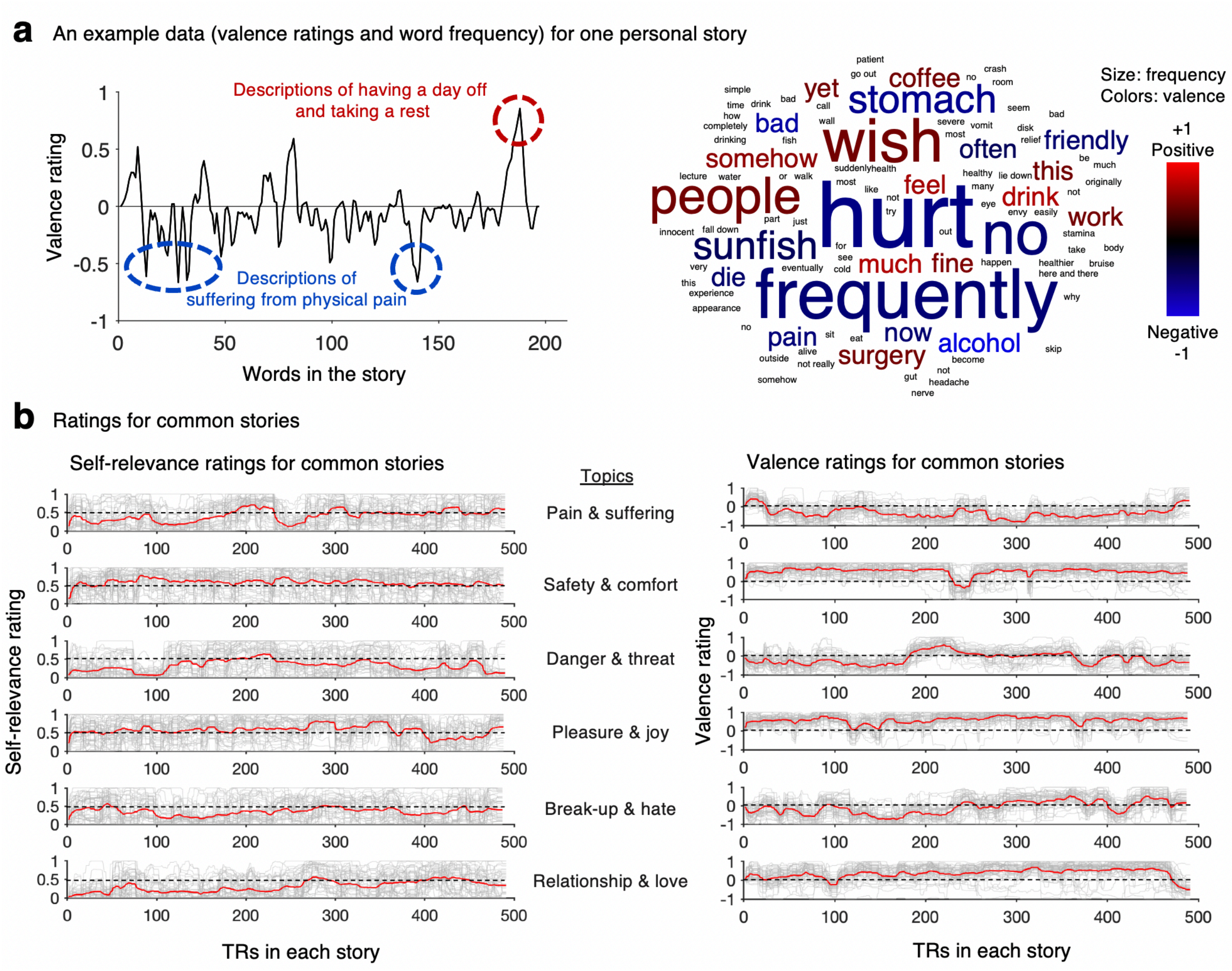


**Supplementary Fig. 1 (related to Fig. 1). Example behavioral data. a,** An example of valence ratings for one personal story. Left: The line plot shows the valence ratings over the words in the story. The topic of the story was pain and suffering. The blue circle areas show the parts in which the participant described suffering from physical pain, and thus the valence ratings were negative. The red circle area is about having a day off and taking a rest, and thus the valence rating for that part was the highest. Right: The word cloud shows the frequency (size) and average valence ratings for each word from the same personal story. The words including 'hurt', 'frequently', 'wish', and 'people' appeared most frequently throughout the story. Blue color indicates the words rated as negative, whereas red color indicates the words rated as positive on average. **b,** Group-average ratings of self-relevance (left) and valence (right) for common stories. There were six common story topics: (1) Pain and suffering, (2) Safety and comfort, (3) Danger and threat, (4) Pleasure and joy, (5) Break-up and hate, and (6) Relationship and love. Each gray line indicates each participant's ratings, and red thick lines indicate group averages.

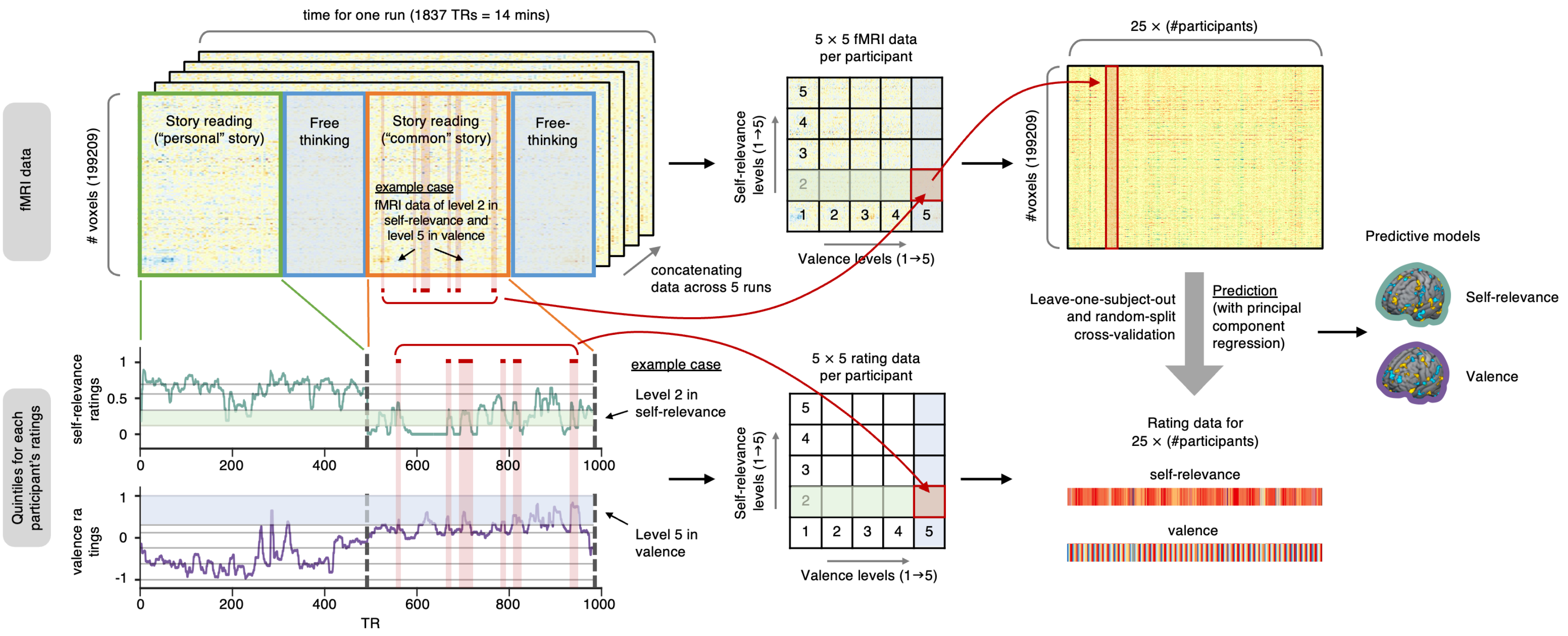


**Supplementary Fig. 2. Predictive modeling analysis pipeline (related to Fig. 2)**. To effectively disentangle fMRI patterns for self-relevance and valence, we manually orthogonalized the data using two-way quantization. First, we divided the data into quintiles (5 levels) separately for self-relevance and valence based on their ratings. We then marked the time points (i.e., TRs) using the levels of two dimensions, resulting in 5×5 data quantization of TR indices. Using these indices (for example, the figure shows level 2 for self-relevance and level 5 for valence as the red-shaded TRs), we averaged the fMRI and rating data, generating 25 fMRI images and rating data per participant. With these orthogonalized data, we trained whole-brain pattern-based predictive models using principal component regression (PCR) with leave-one-subject-out cross-validation (LOSO-CV) and random-split cross-validation (RS-CV).

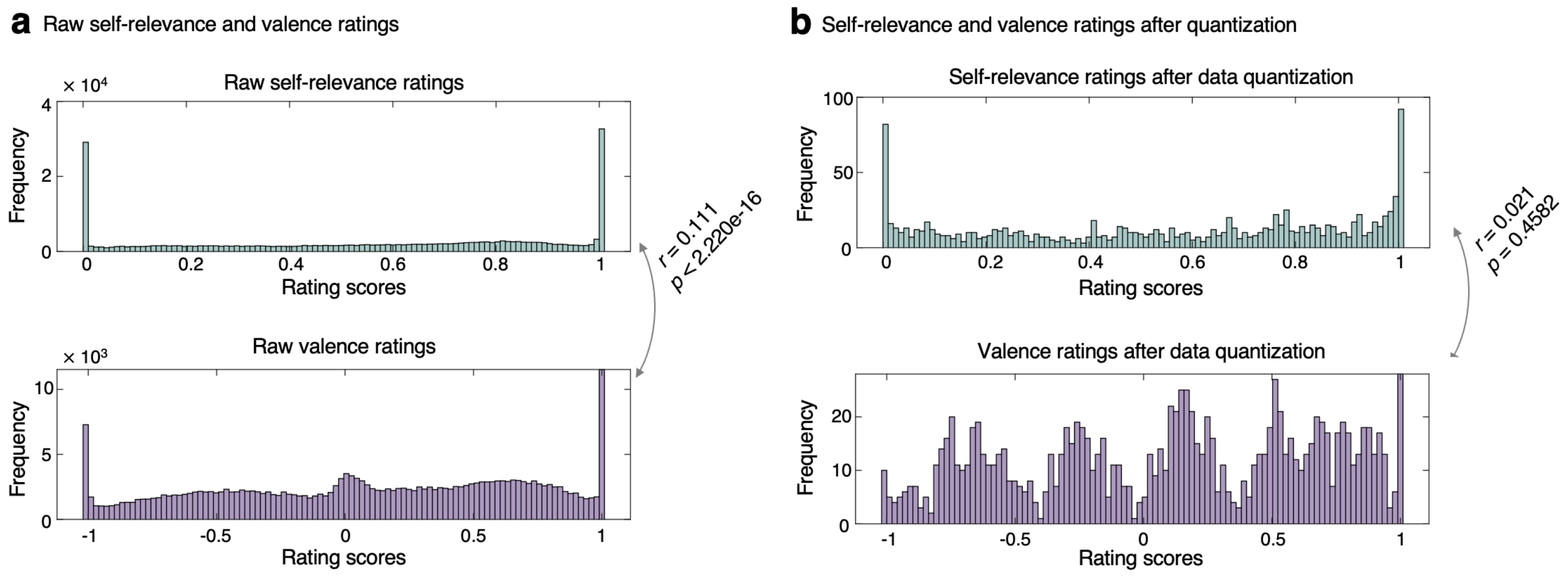


**Supplementary Fig. 3. Distributions of self-relevance and valence ratings before and after data quantization (related to Fig. 2)**. **a**, Distributions of raw self-relevance (upper panel) and valence (bottom panel) ratings at the TR level across all participants. Pearson correlation between raw ratings of self-relevance and valence was $r = 0.111$, $p < 2.220e\text{-}16$. **b**, Distributions of self-relevance and valence ratings after data quantization described in **Supplementary Fig. 2**. Pearson correlation between self-relevance and valence ratings was $r = 0.021$, $p = 0.4582$.

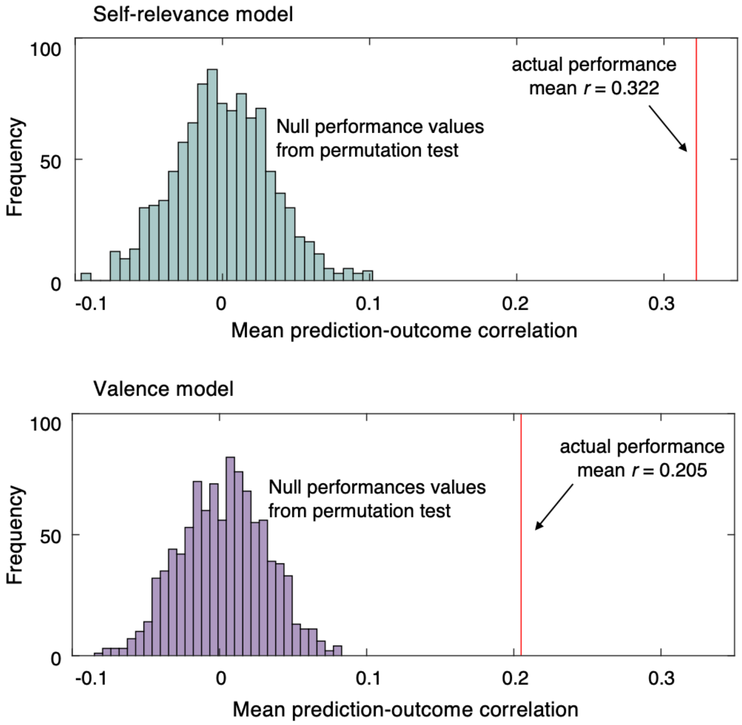


**Supplementary Fig. 4. Results from permutation tests on model performance (related to Fig. 2)**. The histograms (upper panel: self-relevance model, bottom panel: valence model) show the distributions of mean within-participant prediction-outcome correlations from permutation tests with 1,000 iterations. The actual model performance (i.e., mean prediction-outcome correlation) was 0.322 for self-relevance and 0.205 for valence.

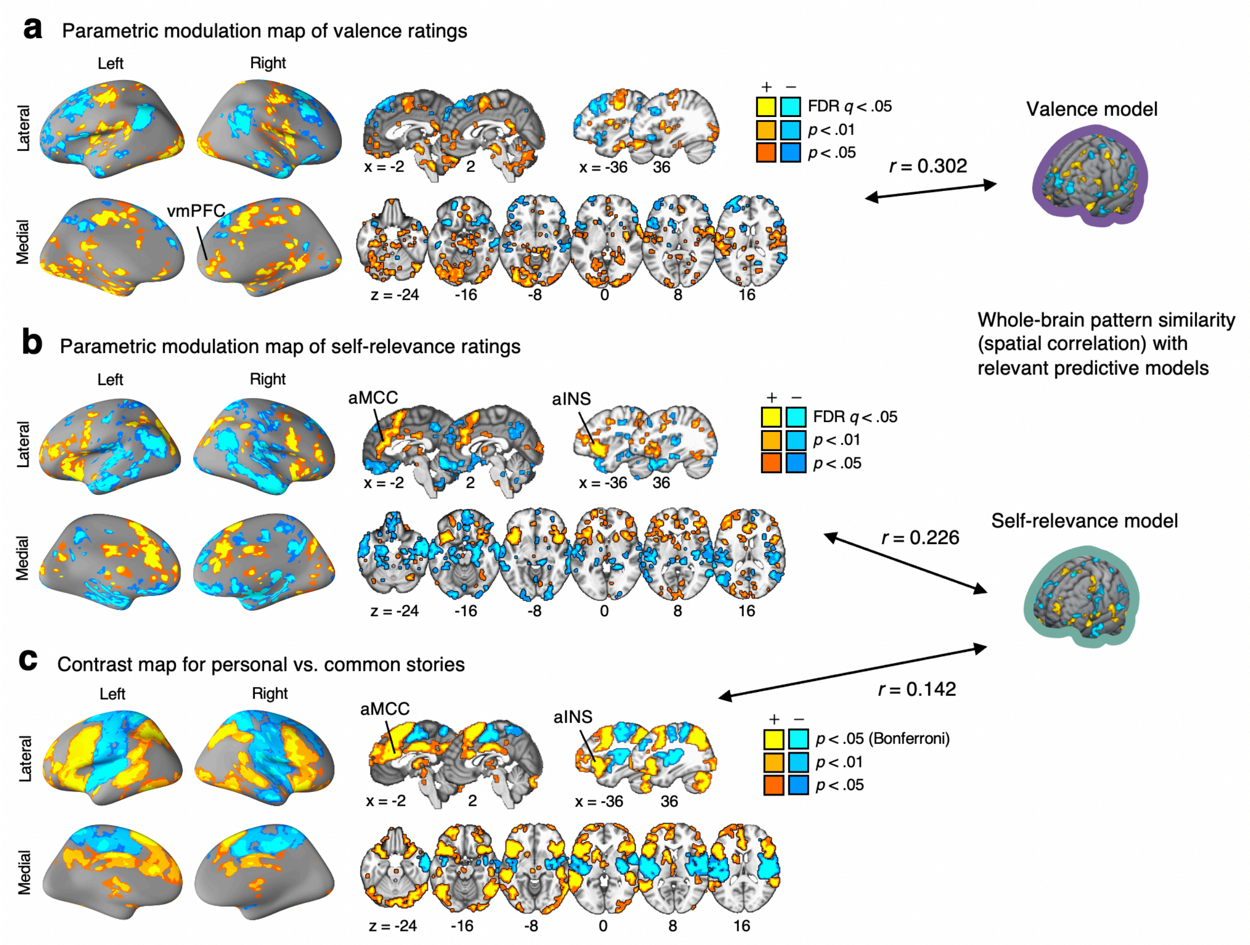


**Supplementary Fig. 5 (related to Fig. 2). General linear model (GLM) results**. **a,** The brain map shows the voxels that are parametrically modulated with the TR-level valence ratings. We thresholded the results with FDR $q < 0.05$ and pruned the map using two more liberal thresholds, uncorrected $p < 0.01$ and $p < 0.05$, two-tailed to show the extent of activation clusters. The brain regions with the positive vs. negative relationship with the valence scores are shown in warm vs. cool colors, respectively. We also show the whole-brain pattern similarity with the predictive model of valence. **b,** The parametric modulation map for self-relevance. **c,** Basic contrast map for personal versus common stories. We thresholded the results at Bonferroni corrected $p < 0.05$ and pruned the map with two additional more liberal thresholds, uncorrected voxel-wise $p < 0.01$ and $p < 0.05$, two-tailed.

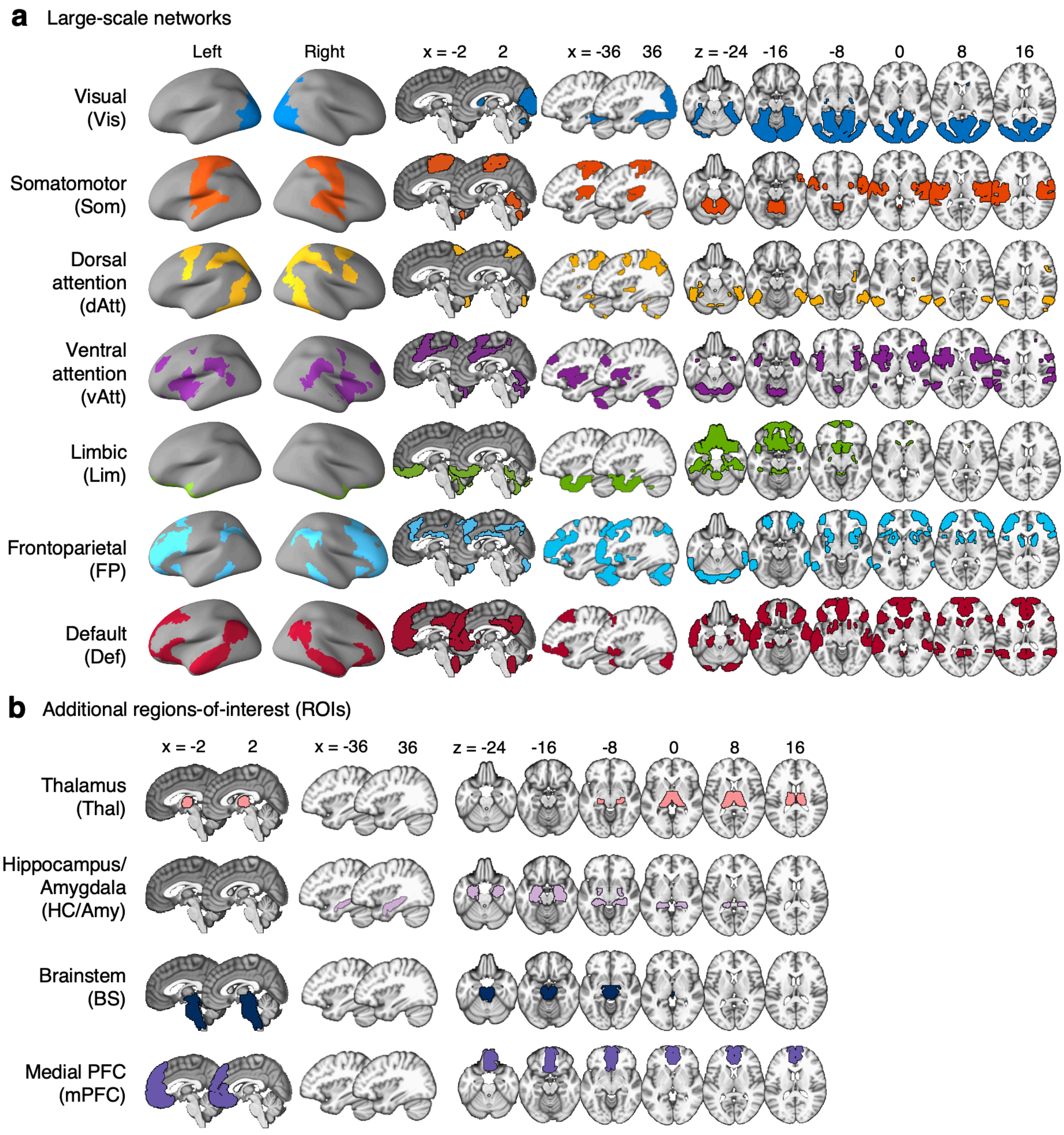


**Supplementary Fig. 6 (related to Fig. 3). Large-scale networks and regions-of-interest.** **a,** Brain maps of the seven large-scale functional brain networks. **b,** Brain maps of the four regions-of-interest (ROIs).

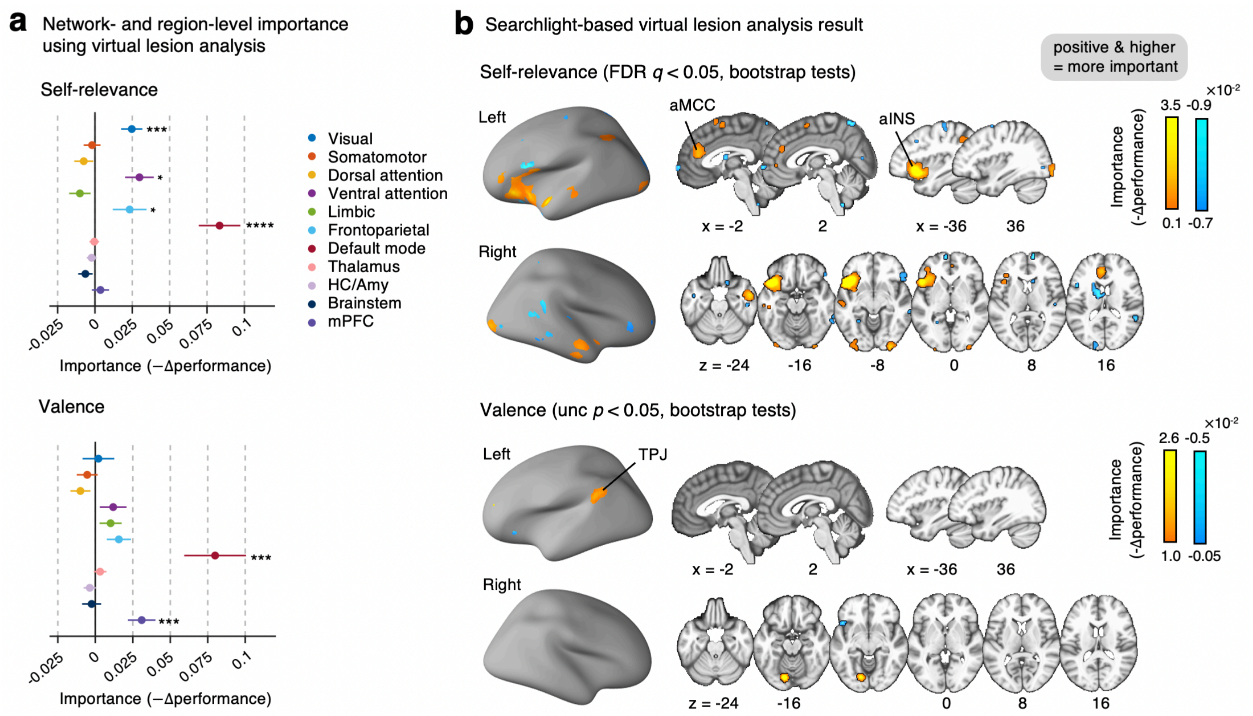


**Supplementary Fig. 7 (related to Fig. 3). Virtual lesion analysis results**. We examined which features were important for predicting self-relevance and valence using the virtual lesion analysis, which calculated the changes in the prediction performance (i.e., prediction-outcome correlation) after excluding a single large-scale network, region, or searchlight at a time. The decreased prediction-outcome correlation after removing the feature indicates that the feature is important for the prediction, while the increased prediction-outcome correlation indicates that the feature is not important. Thus, we defined -Δperformance as the importance of the feature. **a,** The virtual lesion analysis results for the self-relevance model (top) and the valence model (bottom) with the large-scale networks and some ROIs. Each colored dot represents the prediction-outcome correlations for each network or region with bootstrap tests with 10,000 iterations. The error bars represent the standard deviation of the sampling distribution. $^{*}p < 0.05$, $^{***}p < 0.001$, $^{****}p < 0.0001$. **b,** Searchlight-based virtual lesion analysis results for the self-relevance model (top) and the valence model (bottom). The map for the self-relevance model was thresholded at FDR $q < 0.05$, two-tailed, bootstrap tests, and the map for the valence model was thresholded at uncorrected $p < 0.05$. MCC, mid-cingulate cortex; TPJ, temporoparietal junction)

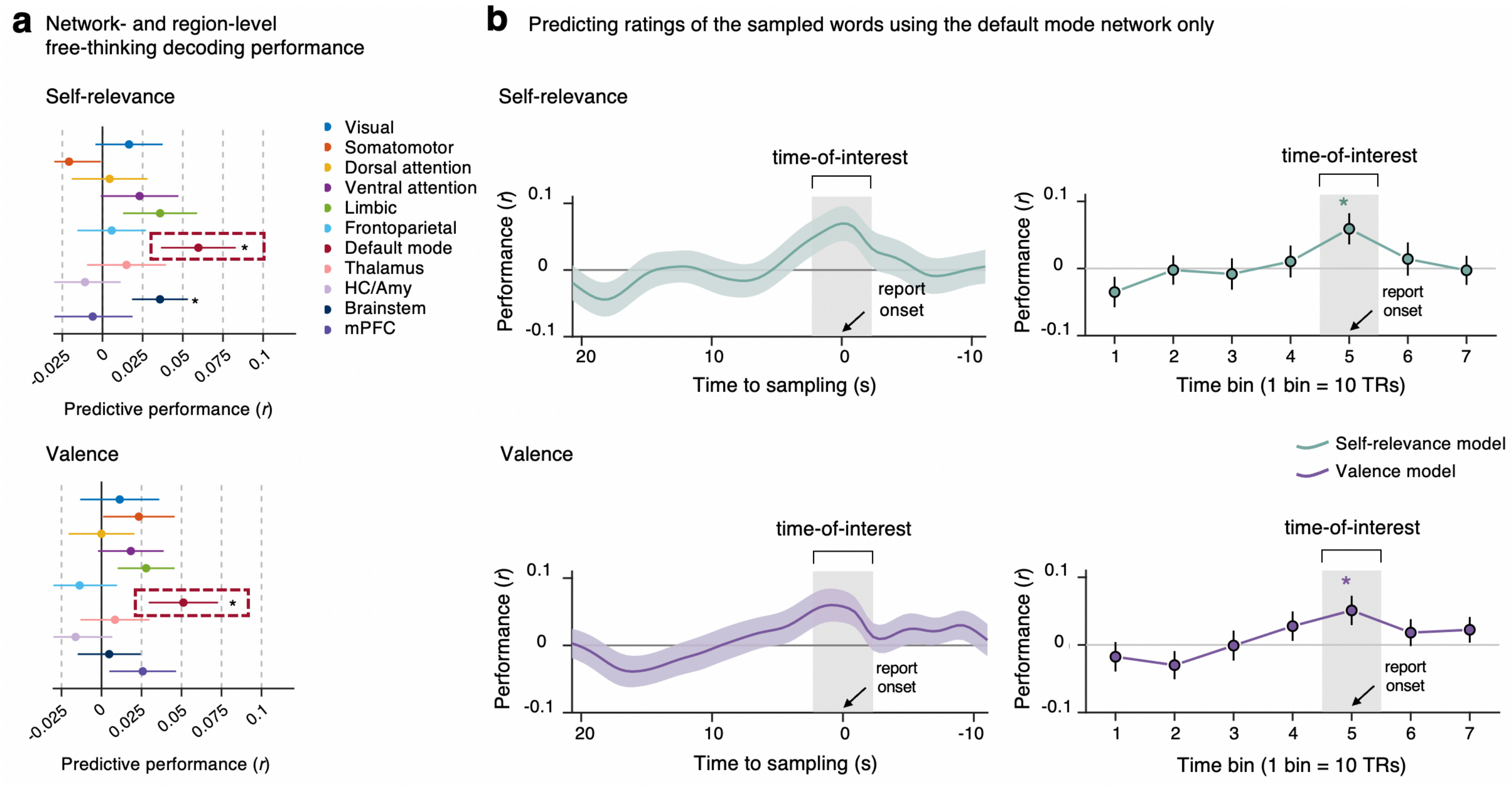


**Supplementary Fig. 8 (related to Fig. 4). Virtual isolation analysis for the decoding of free-thinking. a,** The virtual isolation analysis results for the self-relevance model (top) and the valence model (bottom) with the large-scale networks and some ROIs. Each colored dot represents the prediction-outcome correlations for each network or region with bootstrap tests with 10,000 iterations. The error bars represent the standard deviation of the sampling distribution. **b,** Model performances using the default mode network only (top: predicting self-relevance, bottom: predicting valence) measured by the prediction-outcome correlations. The plots on the left show the prediction performance using a moving-window approach based on the data convolved with the temporal Gaussian kernel (FWHM = 10 TRs). The plots on the right show the prediction performances for the time bins of 10 TRs ($r = 0.0591$ and 0.0510 at the peak for self-relevance and valence, respectively). Shading and error bars indicate the standard deviation of performances across all participants. $^{*}p < 0.05$.

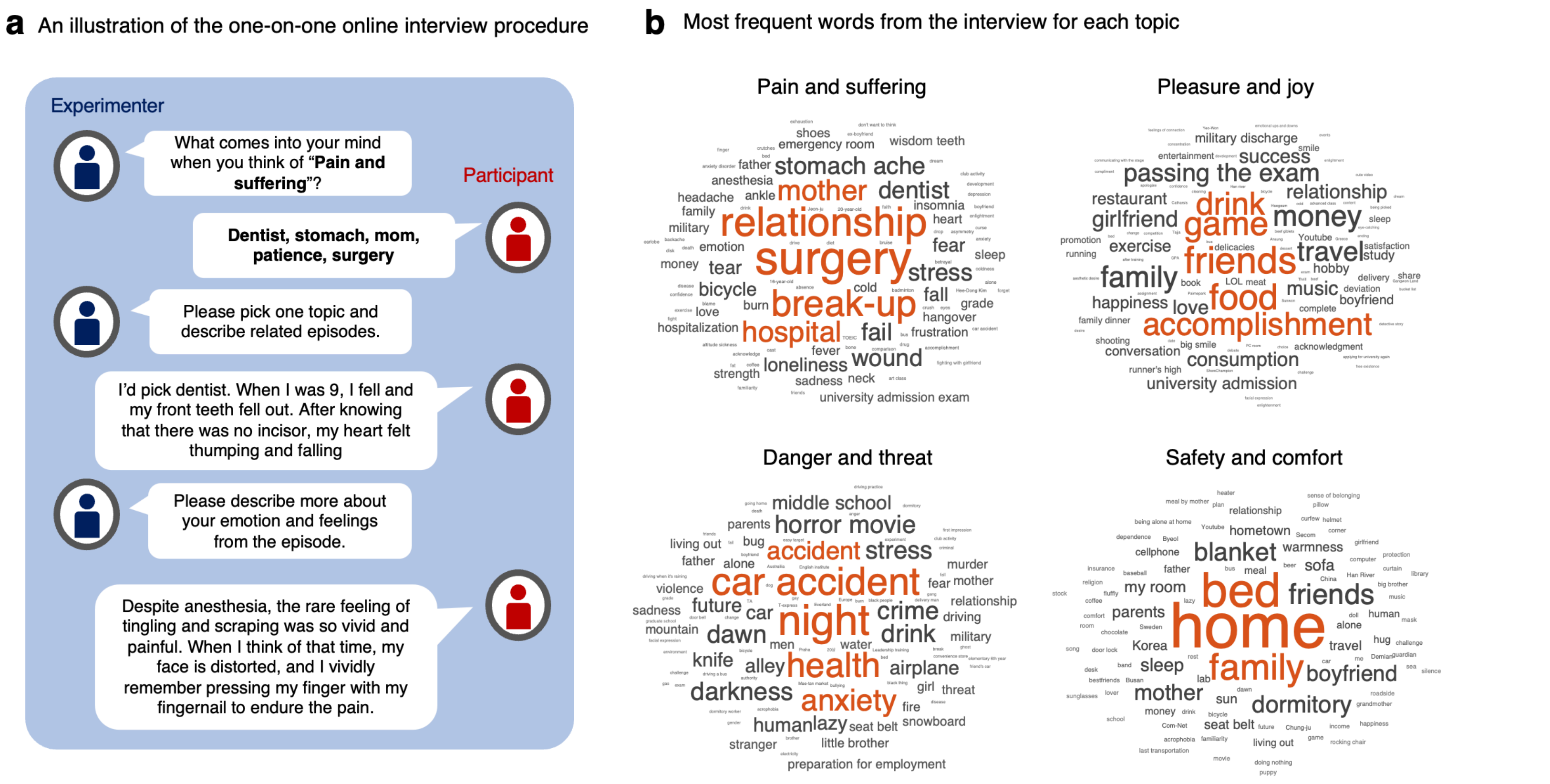


**Supplementary Fig. 9 (related to Fig. 1). a,** An illustration of the one-on-one online interview procedure. Using an online messenger application, we conducted one-on-one interviews with participants to create their personal stories. First, we asked participants to generate some words related to given topics (e.g., pain and suffering, pleasure and joy, danger and threat, and safety and comfort) and share their personal episodes related to the self-generated words (for example, 'dentist' in the figure). We then encouraged participants to share their personal feelings and emotions related to the episode. We asked multiple questions until we gathered content long enough to make a story. **b,** The wordclouds show the words that appeared most frequently for each topic. The size of the words indicates the frequency. For example, for 'pain and suffering,' participants reported words including surgery, relationship, break-up, mother, and hospital frequently.

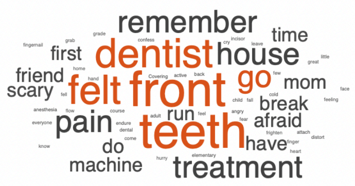


Topic: Pain and suffering
“.. Despite anesthesia, the rare feeling of tingling and scraping was so vivid and painful. So even after a long time, I'm still nervous and frightened..”

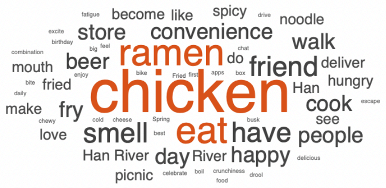


Topic: Pleasure and joy
“.. During this Spring, I ate chicken and ramen after setting up a tent in Han River with friends who celebrated my birthday. Even now, thinking about that day makes me drool. These happy and joyful memories are the driving force for me to live another day.”

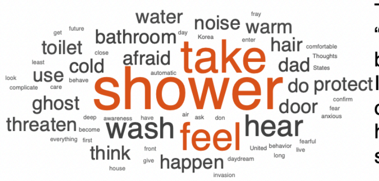


Topic: Danger and threat
“.. The restroom is the most threatened space because I am defenseless. During the shower, I fray my nerve to hear whether the front door opens or closes outside the bathroom. If I hear the sound of a door while taking a shower, I stop everything and look outside. ..”

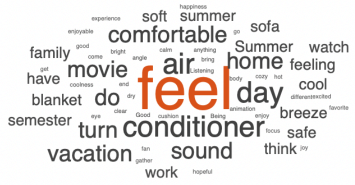


Topic: Safety and comfort
“.. I feel most satisfied than ever inside the cozy, soft blanket while the air conditioner is on. I feel like the whole world is mine when I feel the dry yet cool breeze from the air conditioner while watching my favorite animations and movies on a summer day..”

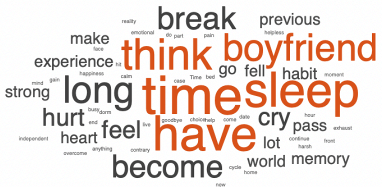


Topic: break-up and hate
“.. He made me wait for about an hour because he was “busy,” and spitted out a goodbye in front of my face. At that time, I came back home crying a lot. I cried, fell asleep, and cried again, then fell asleep — the cycle went on and on...”

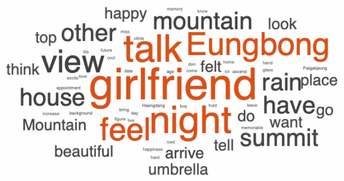


Topic: Relationship and love
“..Eungbong mountain is a dating place I used to go with my ex-girlfriend… As we talked seriously at the summit, our trust in each other increased. I also felt a lot of happiness when I did sincere love. I feel so happy when I think about it now, but I feel sick because I think I will never feel it again”

**Supplementary Fig. 10 (related to Fig. 1). Common stories**. In addition to four personal stories, we made a common set of six stories that were used for all participants. We created the stories through one-on-one interviews with pilot participants using the same four topics (i.e., pain and suffering, pleasure and joy, danger and threat, and safety and comfort) and two additional topics—‘relationships and love’ and ‘break-ups and hatred.’ The word frequency for each common story is illustrated using wordclouds on the left, and some excerpts from each story are shown on the right. All stories were presented in Korean during the experiment.

**Supplementary Table 1. Prediction performance of virtual isolation and lesion analyses (bootstrap tests with 10,000 iterations).**

| Network/regions | | Predicting self-relevance | | | | Predicting valence | | | |
|---|---|---|---|---|---|---|---|---|---|
| | | Virtual isolation analysis | | Virtual lesion analysis | | Virtual isolation analysis | | Virtual lesion analysis | |
| | | Mean *r* | *P* | -Δperformance | *P* | Mean *r* | *P* | -Δperformance | *P* |
| Networks | Vis | **0.1648** | **< 0.0001** | **0.0246** | **< 0.001 (0.0005)** | 0.0336 | 0.2028 | 0.0021 | 0.8447 |
| | Som | 0.0241 | 0.3778 | -0.0019 | 0.7316 | 0.0209 | 0.4081 | -0.0054 | 0.4386 |
| | dAtt | 0.0374 | 0.2663 | -0.0075 | 0.2374 | -0.0269 | 0.3708 | -0.0099 | 0.1352 |
| | vAtt | **0.1720** | **< 0.0001** | **0.0298** | **< 0.05 (0.0019)** | **0.1182** | **< 0.0001** | 0.0119 | 0.1814 |
| | Lim | 0.0281 | 0.4716 | -0.0101 | 0.1613 | **0.0832** | **< 0.01 (0.0031)** | 0.0102 | 0.1668 |
| | FP | **0.1213** | **< 0.0001** | **0.0232** | **< 0.05 (0.0401)** | **0.1029** | **< 0.0001** | 0.0157 | 0.0514 |
| | DMN | **0.2724** | **< 0.0001** | **0.0834** | **< 0.0001** | **0.1785** | **< 0.0001** | **0.0799** | **< 0.001 (0.0001)** |
| Regions | Thal | 0.0221 | 0.4395 | -0.0005 | 0.8854 | 0.0375 | 0.2615 | 0.0032 | 0.4495 |
| | HC/amy | 0.0111 | 0.7274 | -0.0022 | 0.4920 | 0.0031 | 0.9052 | -0.0037 | 0.3676 |
| | BS | -0.0164 | 0.5718 | -0.0064 | 0.1835 | 0.0097 | 0.6856 | -0.0025 | 0.6989 |
| | mPFC | **0.0733** | **< 0.05 (0.0139)** | 0.0037 | 0.5228 | **0.1083** | **< 0.01 (0.0017)** | **0.0309** | **< 0.001 (0.0008)** |

***Note.*** The *r* values for the virtual isolation analysis indicate the prediction-outcome correlation, which was used to evaluate the importance of the network or region in predicting self-relevance and valence. The -Δperformance for the virtual lesion analysis indicates the changes in the prediction performance (i.e., prediction-outcome correlation) after excluding a single large-scale network or region from the model at a time. p values were calculated using bootstrap tests with 10,000 iterations. Vis, visual; Som, somatomotor; dAtt, dorsal attention; vAtt, ventral attention; Lim, Limbic; FP, frontoparietal; DMN, default mode networks; Thal, thalamus; HC, hippocampus; amy, amygdala; BS, brainstem; mPFC, medial prefrontal cortex.
